# Post-correlation on-lamella cryo-CLEM reveals the membrane architecture of lamellar bodies

**DOI:** 10.1101/2020.02.27.966739

**Authors:** Steffen Klein, Benedikt H. Wimmer, Sophie L. Winter, Androniki Kolovou, Vibor Laketa, Petr Chlanda

## Abstract

Lamellar bodies (LBs) are surfactant-rich organelles in alveolar cells. LBs disassemble into a lipid-protein network that reduces surface tension and facilitates gas exchange in the alveolar cavity. Current knowledge of LB architecture is predominantly based on electron microscopy studies using disruptive sample preparation methods. We established and validated a post-correlation on-lamella cryo-correlative light and electron microscopy approach for cryo-FIB milled cells to structurally characterize and validate the identity of LBs in their unperturbed state. Using deconvolution and 3D image registration, we were able to identify fluorescently labeled membrane structures analyzed by cryo-electron tomography. In situ cryo-electron tomography of A549 cells as well as primary Human Small Airway Epithelial Cells revealed that LBs are composed of membrane sheets frequently attached to the limiting membrane through “T”-junctions. We report a so far undescribed outer membrane dome protein complex (OMDP) on the limiting membrane of LBs. Our data suggest that LB biogenesis is driven by parallel membrane sheet import and by the curvature of the limiting membrane to maximize lipid storage capacity.

## Introduction

Lamellar bodies (LBs) are specialized organelles exclusively found in alveolar type 2 epithelial cells (AEC2) and in keratinocytes^1^. Alveolar LBs produce, accumulate and secrete surfactant, a mix of specialized lipids and proteins. Upon secretion into the alveolar cavity, it rapidly disassembles into a highly organized network. Pulmonary surfactant reduces the surface tension at the air-water interface in the alveoli to facilitate gas exchange during respiration. Therefore, it must be constantly replenished to sustain breathing^2^. AEC2 and surfactant are primary players in the pulmonary immune response^3^. Defects in surfactant production are associated with increased risk of respiratory infection by pathogens such as influenza A virus^4^, respiratory syncytial virus^5^, pneumonia^6^ and mycobacterium tuberculosis^7^. Surfactant protein D (SP-D) specifically binds glycosylated pathogens, including SARS-CoV-1^8^. AEC2 fatty acid metabolism and LB ultrastructure are severely disrupted by the pandemic influenza strain H1N1^9^ and the highly pathogenic H7N9 strain^10^. In spite of their importance in health and disease, many questions remain open about LB biogenesis, structure and secretion. LBs are composed of a core containing multilamellar membrane sheets surrounded by a limiting membrane as revealed by thin-section transmission electron microscopy (TEM)^11^. LBs contain 85% phospholipids by weight, mostly dipalmitoyl-phosphatidylcholine (DPPC), unsaturated phosphatidylcholines, phosphatidylglycerols^12^, as well as cholesterol and specialized surfactant proteins A, B and C (SP-A, SP-B, SP-C)^13^. The majority of the LB-associated proteins are commonly found in lysosomes, LBs are thus classified as lysosome-related organelles. Mass spectrometry identified 34 proteins unique to lung LBs^13^. While the core contains the small hydrophobic proteins SP-B and SP-C^14^, the limiting membrane is enriched in the flippase ATP Binding Cassette Subfamily A Member 3 (ABCA3)^15^.

In the current model of LB biogenesis, lipids are flipped by ABCA3 from the cytosolic to the luminal leaflet and are imported into the LB core^16^, where SP-B and SP-C are responsible for further lipid rearrangement into tightly packed membrane sheets^17^. However, this model has been difficult to validate. The LBs’ high lipid content is poorly preserved as a result of room temperature TEM sample preparation, which relies on chemical fixation and dehydration. In consequence, the concentric membranes inside the LB appear wrinkled. Therefore, it is neither understood how they are organized in three-dimensions (3D) nor is it known how the membrane stacks are formed. A study employing cryo-electron microscopy of vitrified sections (CEMOVIS) on rat lungs enabled imaging of frozen-hydrated LBs and showed smooth concentric membranes^18^. However, due to compression artifacts caused by sectioning and lack of compatibility with cryo-electron tomography (cryo-ET)^19^, the study provided only little insight into the complex LB architecture.

Unlike CEMOVIS, cryo-focused ion beam (cryo-FIB) milling enables preparation of thin cellular lamellae of arbitrary thickness with a smooth surface and without compression such that they are compatible with cryo-ET^20,21^. Correlative light and electron microscopy (CLEM) enables unequivocal identification of the targeted compartments and yields structural details^22^. CLEM methods have been adapted to cryo-EM and have successfully been implemented on *in vitro* samples^23,24^ or for cryo-ET performed on whole cells^25–28^. However, a correlation of light microscopy (LM) and electron microscopy (EM) data in a workflow involving cryo-FIB milling is challenging due to the milling geometry and multiple transfers between microscopes: each transfer increases the risk of sample devitrification and ice-contamination. So far, available *in situ* cryo-CLEM workflows involving cryo-FIB milling are aimed at site-specific cryo-FIB milling^29,30^.

Here, we show that precise knowledge of the lamella position in the context of the entire cell determined by cryo-LM after cryo-TEM imaging facilitates accurate mapping of the original LM data to cellular structures on the lamella. In the presented workflow, a 2D correlation is applied to target the region of interest for milling in the X-Y plane of the grid. A second, post-correlation step utilizing LM data acquired after cryo-TEM imaging, deconvolution and 3D correlation is then applied to identify the observed structures corresponding to the position of the lamella not only in X-Y but also in the Z dimension. We show that the latter is essential to increase the correlation precision by computationally removing out-of-lamella fluorescent signals.

We applied the post-correlation on-lamella cryo-CLEM workflow to study LBs within A549 cells, a model for AEC231, that were transiently transfected with ABCA3-eGFP, a well-characterized LB marker15. After both correlation steps, 76% of the ABCA3-eGFP signal corresponded to membrane-bound organelles containing either vesicles or lamellated membranes typical for LBs. In situ cryo-ET allowed us to structurally characterize the membrane organization in ABCA3-eGFP positive LBs without sample preparation artifacts. The LB core shows tightly packed membrane sheets with varying curvature. We found parallel bilayer sheets connected perpendicularly to the limiting membrane via “T”-junctions and concentric bilayer sheets as hallmark structures of LBs. In addition, our work revealed a large outer membrane dome protein (OMDP) on the limiting membrane of some LBs, presumably involved in their formation and trafficking. To corroborate our findings, we analyzed LBs in primary human lung cells, where we observed both “T”-junctions and OMDPs.

## Results

### Design of the post-correlation on-lamella cryo-CLEM workflow

The goal of on-lamella cryo-CLEM is to precisely localize an organelle of interest within the cell both in the X-Y plane and in the Z-dimension. While the former is straightforward, the latter is challenging because (i) the Z-dimension of the lamella is reduced to under 5% of the original Z-dimension of the cell (cryo-lamellae have a thickness of 150 – 200 nm, while an intact cell has a thickness of 4 – 6 µm) and (ii) the cryo-LM has limited Z-resolution. To overcome these difficulties, we implemented a post-correlation on-lamella cryo-CLEM workflow that uses deconvolved cryo-LM 3D maps of the cell before and after cryo-lamella preparation (Fig. 1). We opted for performing the cryo-LM acquisition of the cryo-lamellae after cryo-TEM imaging, as imaging in a cryo-LM increases ice contamination and hence decreases the quality of cryo-ET. This results in a loss of fluorescence on the lamellae due to the electron beam damage occurring during cryo-ET. We therefore use the unimaged surrounding cell body that retains its fluorescence to register the cryo-LM map acquired before and after cryo-TEM acquisition (Supplementary Fig. 1). We use the cryo-LM map acquired after cryo-TEM to identify lamella position and tilt, while we use the initial cryo-LM map acquired before milling to obtain the fluorescence signal. After 3D registration of the two deconvolved cryo-LM maps, it is possible to measure the tilt of the lamella using the brightfield signal in the Z-Y-plane and tilt the complete composite stack accordingly. This enables us to extract a single slice of the registered and tilt-corrected image stack containing only the fluorescent signal corresponding to the lamella. Using rigid transformation, this fluorescence map can finally be correlated to the cryo-TEM map, facilitating the identification of the structure of interest analyzed by cryo-ET. As the correlation of cryo-LM and cryo-TEM maps is performed after cryo-ET, we named this approach “post-correlation on-lamella cryo-CLEM”.

**Figure 1.**
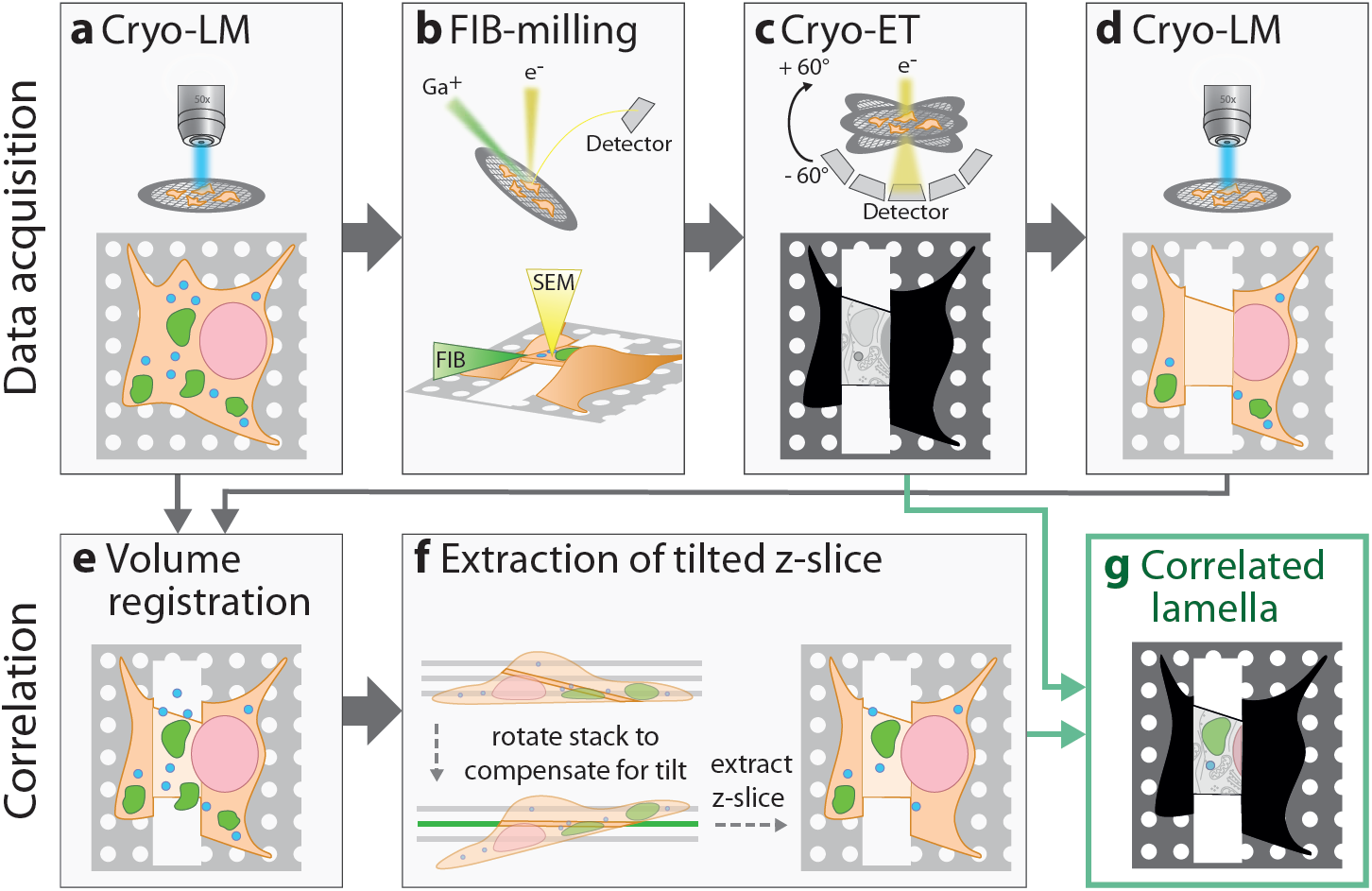
Schematic workflow for post-correlation on-lamella cryo-CLEM. The workflow consists of two parts: data acquisition (a–d) and data processing for correlation analysis (e–g). Structures in cells grown on EM-grids are fluorescently labeled prior to vitrification by plunge-freezing. **a**, In the first step, the vitrified sample is mapped by cryo-LM, acquiring Z-stacks with a 300 nm spacing. **b**, In the second step, labeled cells are thinned by cryo-FIB-milling to produce lamellae of 150 - 200 nm thickness. **c**, Lamellae are mapped by cryo-TEM and tilt series of areas of interest are acquired. **d**, FIB-milled areas are mapped again by cryo-LM to retrieve the exact position of the lamellae. For both Z-stacks of step (a) and step (d), deconvolution is performed to increase the resolution (not shown in the figure). **e**, To combine the information of the fluorescent signal from step (a) and the lamella position from step (d), both Z-stacks are aligned using an automated 3D registration algorithm. After image registration, the transmitted light bright field (TL-BF) channel of step (d) is combined with the fluorescent channels of step (a) leading to a combination of lamella position and fluorescent signal in a single composite Z-stack. **f**, To compensate for lamella tilt, the tilt is measured using the TL-BF signal, the Z-stack is rotated accordingly and a Z-slice corresponding to the lamella is extracted. **g**, Finally, the extracted Z-slice is registered to the cryo-TEM map from step (c).

As a proof of principle, we initially applied the post-correlation on-lamella cryo-CLEM workflow to localize fluorescently labelled lipid droplets (LDs) in A549 cells because LDs are easily distinguishable by cryo-EM. We could validate all LDs (n = 11) identified by cryo-EM on 4 cryo-lamellae by our correlation approach (Supplementary Fig. 2) with an average X-Y correlation precision of 124 nm (standard deviation (SD) = 41 nm, n = 11). We next analyzed the impact of the tilt correction and extraction of a single X-Y slice of our workflow. To this end, we repeated the correlation of the 4 lamellae, but instead of an extracted X-Y slice of the fluorescent map, we used a maximum intensity projection (MIP). The comparison of both correlations (Supplementary Fig. 3) revealed that the tilt correction and extraction of a single X-Y slice is essential for the workflow and reduces the out-of-lamella signal by 66%. The correlation using a MIP fluorescent map showed 35 out-of-lamella LD signals which correspond to LDs removed during cryo-FIB milling. By tilt correction and extraction of a single X-Y slice we were able to reduce the number from 35 to 12 out-of-lamella signals.

### Post-correlation on-lamella cryo-CLEM workflow shows ABCA3-eGFP correlation predominantly correlates with lamellated membranous organelles

To localize LBs in A549 cells, we used ABCA3-eGFP overexpression, which induces the formation of LB-like organelles^16^. At 48 h post-transfection, A549 cells contained a median of 127 large spherical structures per cell (SD = 50, range 45 – 179). The ABCA3-eGFP signal predominantly localized to the limiting membrane and exhibited an average diameter of 1.2 µm (SD = 0.8 µm, range 0.1 µm – 4.5 µm) as revealed by confocal microscopy (Supplementary Fig. 4). These measurements are in line with the previously reported LB diameter (range 0.1 – 2.4 µm), based on EM studies performed on lung cells^1^. We transiently transfected A549 cells grown on gold EM-grids and additionally stained them with a nucleus dye and a neutral lipid dye labeling LDs with the intention to use them as fiducial markers, thereby increasing the correlation precision. However, ABCA3-eGFP overexpression leads to a depletion of LDs in the cells. In addition, the neutral lipid dye is also localized to the core of ABCA3-eGFP positive organelles (Supplementary Fig. 5) preventing us from using them as fiducial markers. After vitrification by plunge freezing, the sample was transferred under cryogenic temperatures to a cryo-LM to map the EM-grid. Z-stacks in transmitted light brightfield (TL-BF) and fluorescent channels were acquired covering a large area (ca. 1.2 mm × 1.2 mm) of the grid to maximize the number of potential areas for subsequent milling. To improve contrast and reduce the signal of stray light of the wide-field microscope setup, as well as to improve the correlation precision, we performed deconvolution of the cryo-LM data (Supplementary Fig. 6). After deconvolution, we were able to discriminate closely apposed ABCA3-eGFP positive organelles, which prior to deconvolution appeared as one organelle (Supplementary Fig. 6d-j). Maximum intensity projection (MIP) and stitching of cryo-LM data were used to generate a map, from which suitable areas exhibiting large ABCA3-eGFP positive organelles were selected for milling (Fig. 2a). In most cases, stress relief cuts positioned next to the cryo-lamellae were used to reduce bending of cryo-lamellae before the final milling step^32^. Grids containing 4 – 6 self-supporting lamellae (Fig. 2b) were transferred to a cryo-TEM where they were mapped to judge overall lamella quality and to localize membrane-bound organelles (Fig. 2c). Among frequently present organelles such as mitochondria and nuclei, membrane-bound structures with a lamellated membrane architecture were selected for cryo-ET as putative LBs. To obtain a fluorescent map of the milled cell for on-lamella post-correlation, grids were mapped a second time by cryo-LM after retrieval from the cryo-TEM (Fig. 2d). Milled areas on the grid did not show any specific fluorescent signal, likely due to electron-induced beam-damage, whereas the fluorescence in the surrounding body of the cell was preserved (Supplementary Fig. 1). In some cases, ice contamination detected in TL-BF on the lamella resulted in an auto-fluorescent signal in all fluorescent channels (Fig 2d, asterisk). Fluorescent images of the cell before and after cryo-TEM for all analyzed lamellae are shown in Supplementary Fig. 1. To retrieve the fluorescent signal corresponding to the lamella position we utilized 3D image registration and rigid transformation using one of the deconvolved fluorescence channels to overlay both image volumes (Fig. 2e). The loss of fluorescent signal by using stress relief cuts was negligible and hence did not impede the correlation. The resulting combined image stack incorporates the fluorescent signal before FIB-milling with the TL-BF signal after milling, which contains the precise lamella position. A fluorescent map corresponding to the lamella was extracted from the combined image stack by correcting for the milling angle (Fig. 2f) and finally by extracting the slice corresponding to the plane of the lamella (Fig. 2g). We used the four corners of the lamella as well as clearly identifiable fluorescently labeled markers like the nuclear envelope as landmarks to correlate the extracted Z-slice with the cryo-TEM map (Fig. 2h) by non-rigid transformation. Due to the loss of LDs by ABCA3 overexpression (Supplementary Fig. 5) we were not able to use LDs as fiducial markers which could have further improved the correlation precision in X-Y. A detailed view of the correlated cryo-TEM map showed that the ABCA3-eGFP signal overlaps with multi-lamellated structures (Fig. 2i).

**Figure 2.**
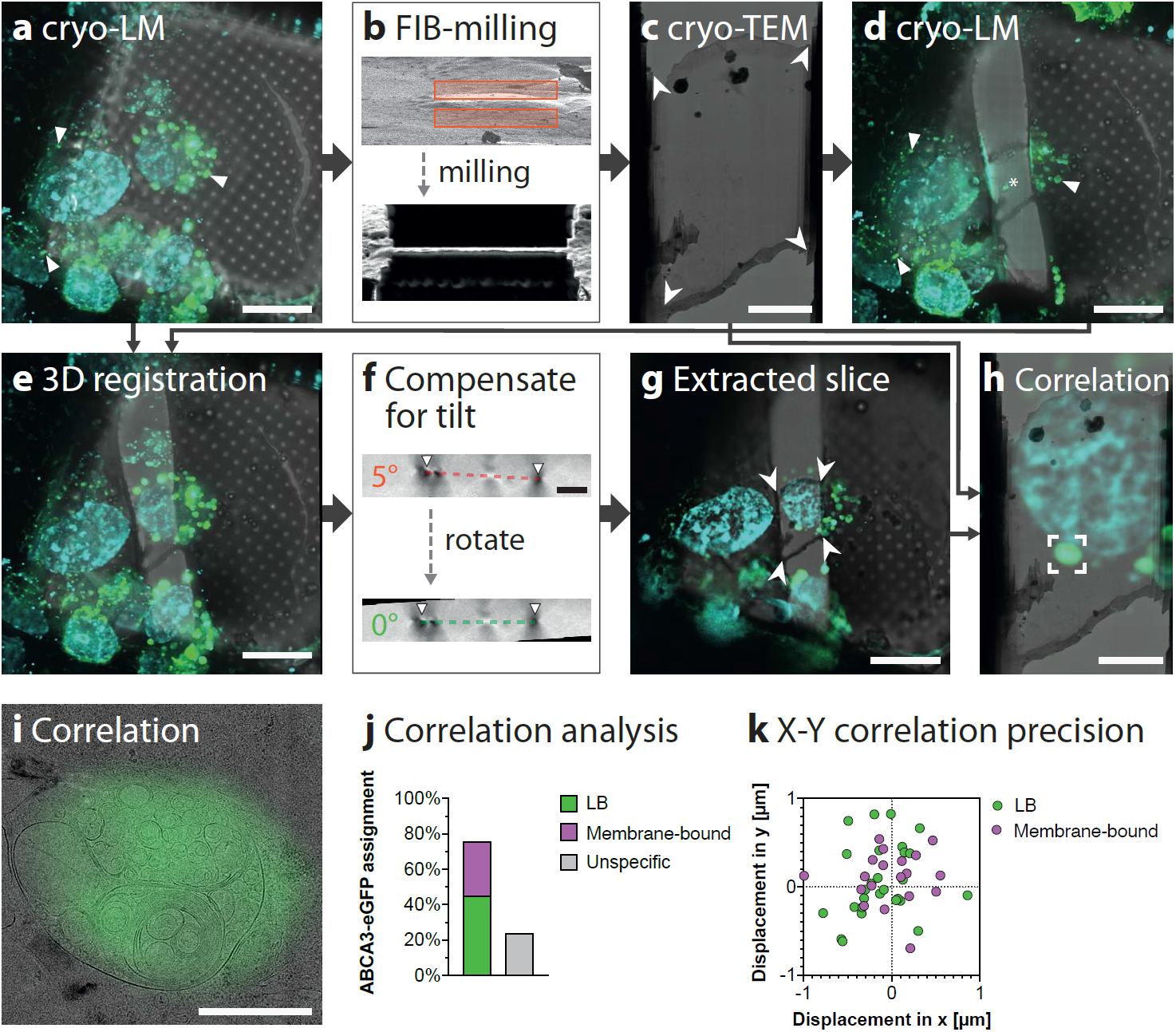
Post-correlation on-lamella cryo-CLEM allows the identification of LB-like structures. A549 cells grown on EM-grids were transiently transfected with ABCA3-eGFP and nuclei and lipid droplets were fluorescently labeled prior to plunge freezing. **a**, MIP of the deconvolved cryo-LM stack acquired before FIB-milling. ABCA3-eGFP is shown in green, and nuclei and lipid droplets are shown in cyan. **b**, Angled FIB image showing the same region before and after milling, respectively. **c**, Stitched cryo-TEM map of the lamella produced by FIB milling acquired at 8,700× magnification. White arrowheads show landmarks used for correlation. **d**, MIP of the deconvolved cryo-LM stack acquired after TEM imaging. The fluorescent signal in the cell body remains visible (arrows). Ice contamination on the lamella (asterisk) shows autofluorescence which is detectable in all fluorescent channels and thus is not emitted from ABCA3-eGFP. **e**, MIP of the registered and aligned image stack. **f**, Z-Y slice showing the lamella tilt corresponding to the FIB milling angle (top) and Z-Y slice after rotation of the Z-stack by 5°. The organometallic platinum coated edges of the lamella (arrows) can be seen as dark dots in the Z-Y-slice. **g**, Extracted Z-slice corresponding to the plane of the lamella with computationally removed the out-of-lamella fluorescent signal. White arrowheads indicate landmarks overlaid with those in (c). **h**, Overlay showing the product of the non-rigid alignment between the TEM map in panel (c) and the fluorescence image in panel (g). **i**, High magnification image of the correlated lamella. **j**, Quantification of correlation with membrane bound organelles. A total of 62 distinct correlated eGFP signals of 6 lamellae were assigned to corresponding structures of the cryo-TEM map. 28 ABCA3-eGFP signals were assigned to LBs, 19 to membrane-bound organelles and 15 could not be assigned to any structure. **k**, Quantification of the correlation precision. For the 47 successfully correlated structures, the distance and angle to the center of the corresponding ABCA3-eGFP signal was measured and shown in a radar plot. The average correlation precision is 450 nm (SD = 249 nm). Scale bars: (a, d, e, g) 20 µm, (c, f, h) 5 µm, (i) 1 µm.

We classified structures corresponding to ABCA3-eGFP isolated signals (n = 62) on 6 lamellae. As a result, 45% of ABCA3-eGFP signal could be assigned to organelles containing lamellated membranes with a displacement smaller than 1 µm. 31% of the ABCA3-eGFP signal was correlated to membrane-bound structures including multivesicular bodies (MVBs), endoplasmic reticulum (ER) or vesicles. The remaining 24% of the ABCA3-eGFP signal could not be assigned to any membrane-bound organelles related to ABCA3 (Fig. 2j, Supplementary Table 1). Correlated structures showed an average displacement to the local maxima of the corresponding ABCA3-eGPF signal of 450 nm (SD = 249 nm) (Fig. 2k, Supplementary Table 1). To evaluate the significance of extracting only the signal corresponding to the lamella on this sample, we also performed a correlation based on the MIP of the tilt-corrected assembled stack for 3 lamellae. In contrast to extracted slices, 57% of cryo-LM signals could not be assigned to any membrane-bound organelles observed by cryo-TEM when MIP was used for the correlation. This validates our analysis on LDs (Supplementary Fig 3), showing that tilt correction and the extraction of a single X-Y slice improves the correlation. In summary, we could correlate 76% of ABCA3-eGFP signal with an average X-Y correlation precision of 450 nm to membrane bound organelles from which 45% showed lamellated morphology.

### LBs contain tightly packed parallel-curved membrane sheets frequently connected to the limiting membrane by “T”-junctions

We analyzed the 3D membrane architecture in 11 tomograms containing ABCA3-eGFP positive organelles. All correlated organelles showed intraluminal vesicles and tightly packed membrane sheets of varying curvature and with regular spacing in the core (Supplementary Fig. 7). In LBs showing large concentric lamellated architecture (Fig. 3, Supplementary Movie 1), we observed that the curvature of membrane sheets increases towards the center, concurrent with a spherical 3D organization of the parallel sheets. Detailed analysis of the regular membrane spacing by Fast Fourier transform (FFT) of a central region (Fig. 3b) revealed three major frequencies which correspond to intra-headgroup-distance (3.7 nm^-1^), bilayer width (5.6 nm^-1^) and bilayer repeat (11 nm^-1^). This is supported by a density line plot of the same area (Fig. 3d). In addition, we observed a crystalline lipid structure with a lateral repeat of 3.4 nm as measured by FFT (Fig. 3g).

**Figure 3.**
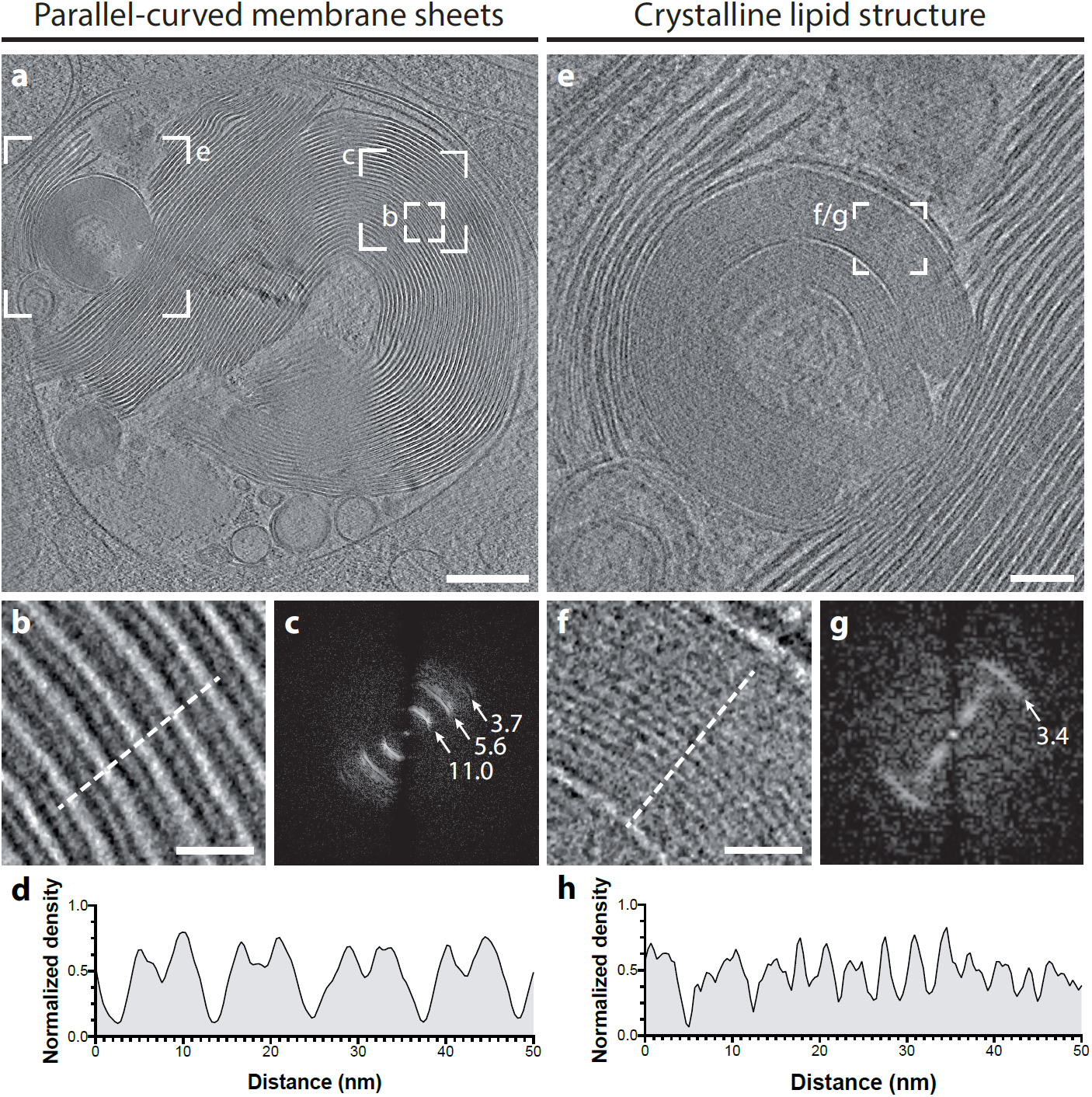
Cryo-ET of LB organelles reveals parallel-curved membrane packaging of membrane sheets and crystalline lipid structures. a, Central slice of the reconstructed tomogram acquired at the site of the correlated ABCA3-eGFP signal (correlation shown in Supplementary Fig. 7k, l). b, Detailed view of the parallel-curved membrane organization of the LB core. c, FFT analysis of parallel-curved membrane spacing, area used for FFT analysis is indicated in (a). FFT analysis reveals three major frequencies at 3.7 nm^-1^ (headgroup-headgroup), 5.5 nm^-1^ (bilayer width) and 11 nm^-1^ (bilayer repeat). d, Plot of a line profile indicated by a dotted line in (b) shows membrane spacing. e, Detailed view of crystalline lipid structures inside LB. f, Parallel sheets observed in the crystalline lipidic core. g, FFT analysis of parallel sheets of crystalline lipid structure, area used for FFT analysis is indicated in (e). FFT analysis reveals a frequency of 3.4^-1^ nm for the spacing. h, Plot of a line profile indicated by a dotted line in (f). Reconstructed tomogram is visualized as Supplementary Movie 1. Scale bars: (a) 200 nm, (e) 50 nm, (b, f) 50 nm.

Although membrane sheets in parallel-curved arrangements were most frequently found in LBs, spiral-coiled membrane sheets, as well as membranes forming closed compartments were also observed within the core of the LB (Fig. 4, Supplementary Movie 2 and 3). Since cellular membranes predominately form closed compartments, we next analyzed the ends of the parallel membrane sheets to provide insight on how the bilayer terminus is structurally organized. Rounded densities at the bilayer termini were frequently observed indicating that the acyl chains of the phospholipids are not exposed to the aqueous surrounding (Fig. 4c–e, arrows). Interestingly, membrane sheets were typically found perpendicularly oriented towards the LB limiting membrane. These sheets were often connected to the limiting LB membrane with a thin density (Fig. 4c). Based on the perpendicular shape of the connection, we call these direct contacts “T”-junctions.

**Figure 4.**
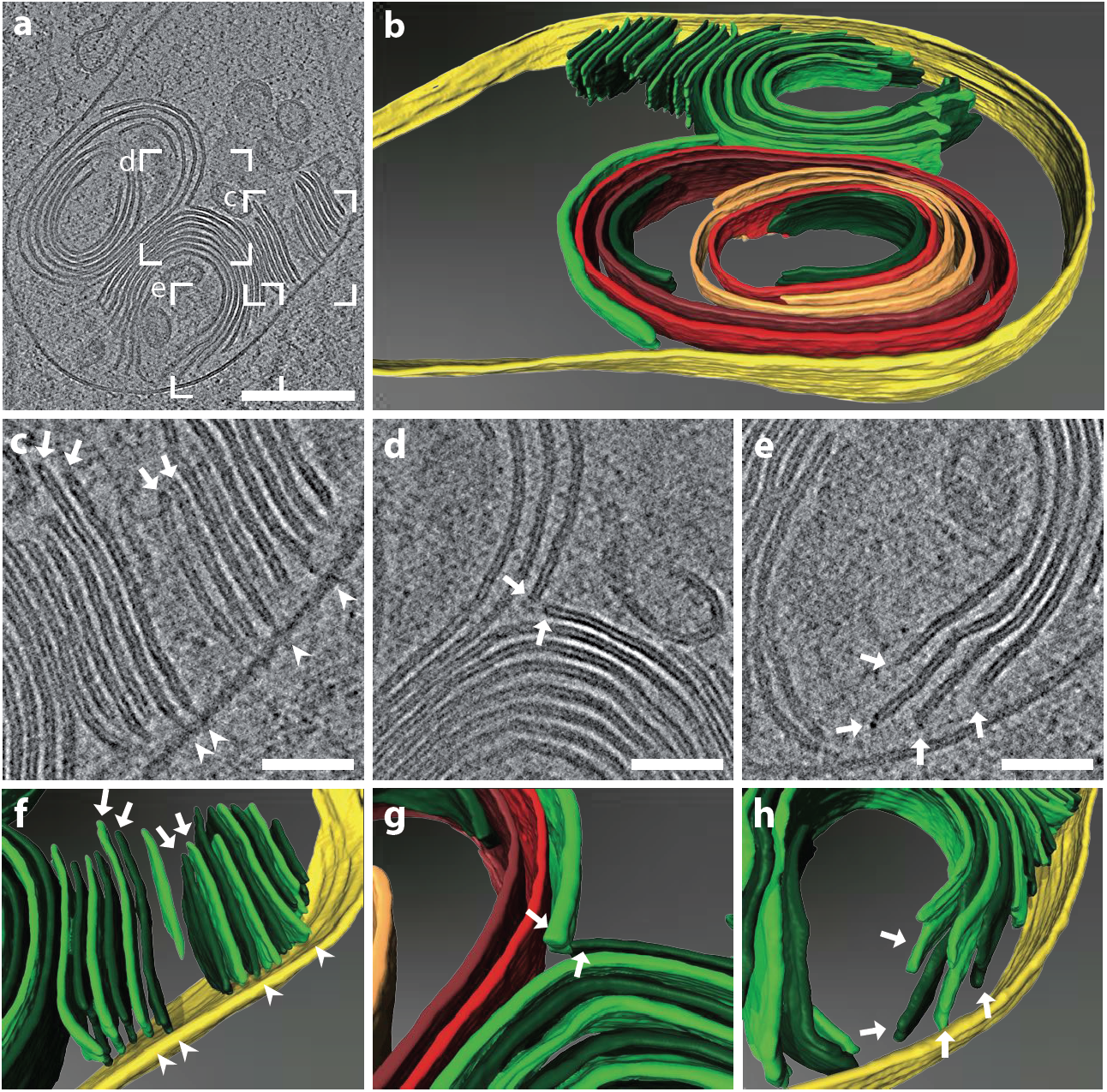
Cryo-ET of LB organelles reveals structural details of membrane sheet termini and “T”-junctions. **a**, Central slice of the reconstructed tomogram acquired at the site of the correlated ABCA3-eGFP signal (correlation shown in Supplementary Fig. 7a, b). **b**, Manual rendering of the tomogram (a). The limiting membrane of the LB is labelled in yellow, membrane sheets are labelled in green. To distinguish individual membrane sheets, alternating shades of green were used. Continuous membranes are labelled in red. A spiral-coiled membrane sheet is labelled in orange. **c–e**, Detailed views of reconstructed tomogram. Perpendicularly oriented membrane sheets towards the limiting membrane are often connected via a thin density (“T”-junction) to the limiting membrane of the LB (arrowheads). Membrane sheets show a rounded density at the membrane termini (arrows). **f–g**, 3D rendering of detailed views (c–e). This figure is visualized as Supplementary Movie 2 and 3. Scale bars: (a) 200 nm, (c, d, e) 50 nm.

### Dome-shaped protein complexes localize at the limiting membrane of LBs

After closer inspection of LBs’ limiting membranes, we observed a dome-shaped protein complex, hereafter referred to as outer membrane dome protein (OMDP) (Fig. 5). OMDPs appeared hollow and were found in the proximity of ER cisternae partially decorated with ribosomes (Fig. 5a–d). Although OMDPs were rare (n = 4) in ABCA3-eGFP transfected A549 cells, they were frequently observed (n = 171) on the limiting membrane of LBs and MVBs in A549 long-term culture from 9 tomograms (Supplementary Fig. 9) as well as on the limiting membrane of LBs in HSAEpC cells (n = 4) (Fig. 5e–h). To obtain insights into the OMDP structure, we manually extracted subtomograms (n = 105) containing OMDPs from 2 tomograms and performed subtomogram averaging (STA) in Dynamo^33^. The STA of OMDP revealed a hollow dome-shaped structure (Fig. 5i–l) with a base diameter of 30 nm, height of 22 nm with a kink at 7 nm and top diameter of 16 nm. Symmetry search performed on the calculated average indicated that OMDP has a C_8_ symmetry.

**Figure 5.**
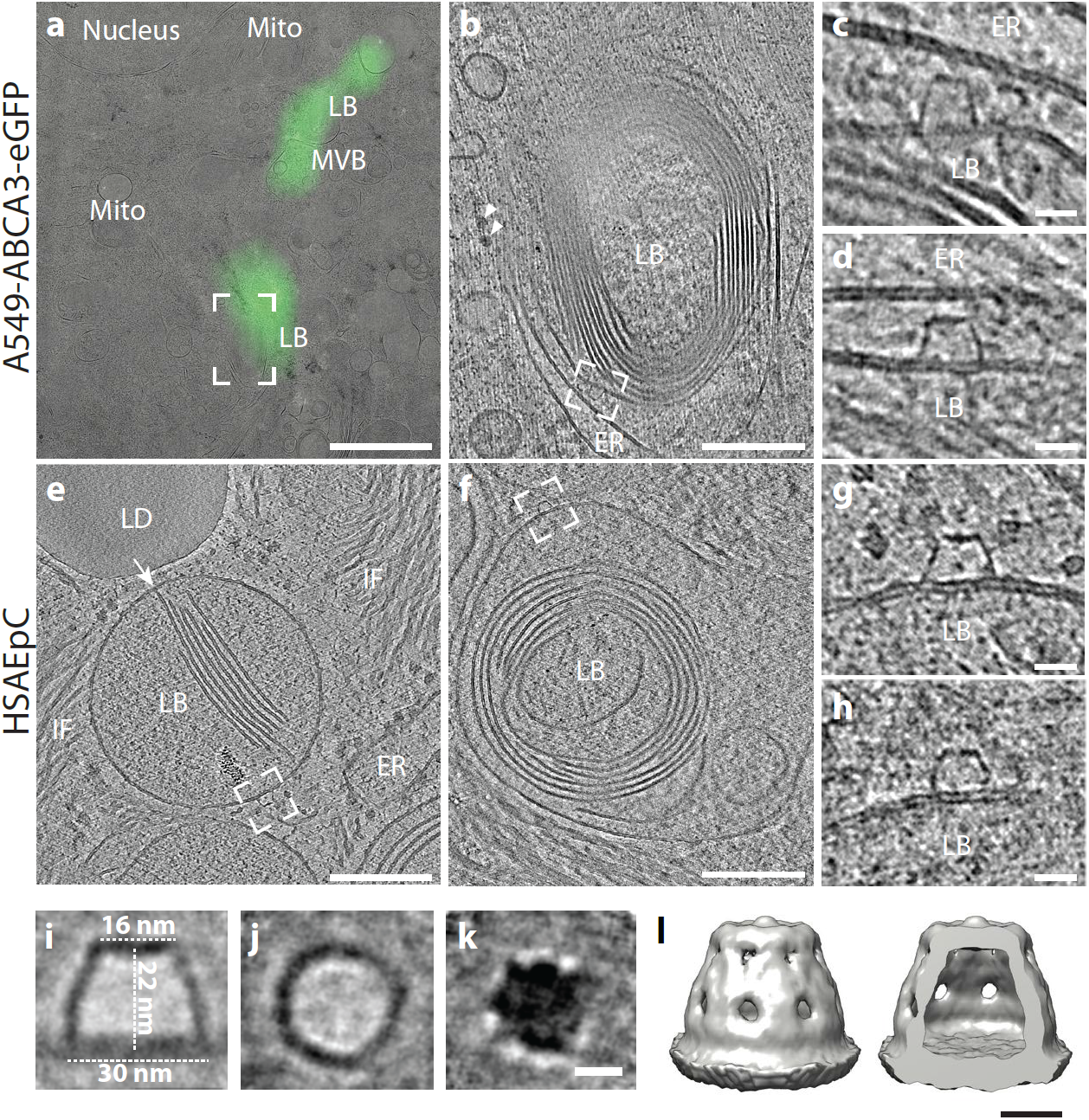
Outer membrane dome protein (OMDP) observed on the limiting membrane of LBs in A549-ABCA3-eGFP and HSAEpC primary lung cells. a, Correlated image of a lamella with ABCA3-eGFP signal overlapping with a MVB and lamellar bodies. Nucleus and mitochondria (Mito) do not show the ABCA3-eGFP signal. b, Slice of a tomogram corresponding to the region indicated in (a) The ER cisterna is indicated. Ribosomes are marked with arrowheads. c, d, Two different slices of the same tomogram corresponding to the area indicated in (b). OMDP assembly on a LB limiting membrane facing the membrane of a neighboring ER membrane. In total 3 identical assemblies are present in the tomogram shown in (b). e, Slice of a tomogram showing a LB precursor containing parallel membrane sheets connected to the limiting membrane (arrow) in HSAEpC primary lung cells. The LB is located in the vicinity of intermediate filaments (IF), lipid droplet (LD) and the endoplasmic reticulum (ER). An OMDP is located on the limiting membrane of the LB (dashed square). f, Slice of a tomogram showing a LB with curved membrane sheets and an OMDP located on the limiting membrane of the LB (dashed squares). g, h, Details of a tomogram showing OMDP corresponding to a marked region in (e, f). i—k, Slices through a STA of 105 OMDPs found in 2 tomograms. (i) represents a XZ-slice through the center of the OMDP, (j) shows a XY-slice through the center and (k) through the top of the OMDP. l, Isosurface of the STA after applying C8 symmetry. Scale bars: (a) 5 µm, (b, e, f) 200 nm, (c, d, g, h) 20 nm, (i, j, k) 10 nm, (l) 10 nm.

### Primary lung cells show LB organelles with “T”-junctions and OMDPs

To validate that the architecture of LBs induced by ABCA3-eGFP expression in A549 cells represents the architecture of physiologically formed LBs, we used primary Human Small Airway Epithelial Cells (HSAEpC) isolated from the distal portion of the human respiratory tract, and also included long-term A549 cultures, which were previously shown to recover a differentiated phenotype with enhanced LB production as found in primary AEC2 cell^34^. *In situ* cryo-ET of milled HSAEpC revealed lamellated organelles (Fig 5e–h, Supplementary Fig. 8) in 10 analyzed tomograms consistent with LBs observed in A549 cells overexpressing ABCA3. Importantly, LBs found in HSAEpC contained membrane sheets attached to the LB limiting membrane via T-junctions (Fig 5e) and the curved lamellated membranes were less condensed (Fig 5f). LBs in the HSAEpC cells were frequently found in the proximity to intermediate filaments (Fig 5e). In addition, whole-cell cryo-ET of long-term A549 cells also revealed LBs (Supplementary Fig. 9) in 6 out of 21 analyzed tomograms. They exhibited similar morphologies to LBs produced by ABCA3-eGFP overexpression and had an average diameter of 0.58 µm (SD = 0.25 µm, range 0.24 – 1.1 µm). Consistent with previously published data^34^, we found lamellated structures in only 1 out of 15 analyzed tomograms of short-term non-transfected A549 cells, indicating that frequent splitting of A549 cells is impeding LB formation.

## Discussion

In this study, we present a workflow for post-correlation on-lamella cryo-CLEM and employ it to study the 3D ultrastructure of LBs in unperturbed conditions. CLEM approaches are extensively used with chemically fixed or high-pressure frozen-freeze substituted (HPF-FS) samples to identify target structures and rare cellular events^22^. However, such methods are inevitably compromised by sample preparation artifacts from dehydration, post-staining and sample sectioning. In particular, the structure of membranes and lipid-rich organelles is poorly retained due to lipid extraction occurring during sample dehydration by organic solvents. Cryo-EM allows direct visualization of membranous structures close to their native state. Correlative approaches such as cryo-CLEM can be applied to identify molecules within the vitreous sample under the condition that the sample remains vitrified and free of ice contamination during the transfer between cryo-LM and cryo-TEM^23^. Until now, cryo-CLEM workflows involving cryo-FIB milling rely on high-precision 3D correlation for site-specific milling to target the region of interest. A publication by Arnold *et al*.^*29*^ reported lamella preparation with 3D targeting based on cryo-LM and cryo-FIB/SEM correlation. This workflow was established on suspended cells mixed with fluorescent beads and required a custom-built cryo-stage for a spinning disk confocal microscope. In such an approach, the correlation accuracy is sensitive to the number and distribution of large fiducial beads and drift occurring during the milling^29^. Recently, a prototype microscope with a cryo-LM integrated into the cryo-FIB microscope chamber was developed^35^. This allows monitoring the fluorescent signal during milling and on the finished lamella. This signal can then be directly correlated without a sample transfer.

In this study, we implemented an alternative strategy and developed a robust on-lamella cryo-CLEM approach, taking advantage of post-correlation. It is suitable for adherent cells cultured directly on EM grids without the introduction of fiducial beads, utilizing a commercially available Leica cryo-LM. Although tilt series cannot be acquired based on *a priori* acquired fluorescence information, the post-correlation approach gives the opportunity to correlate only vitreous, high-quality lamellae judged by cryo-TEM. Most of our attempts to perform cryo-LM after milling, but prior to cryo-TEM, resulted in increased ice contamination on the lamellae, thus preventing high-quality cryo-ET. In addition, we rarely detected fluorescence directly on lamellae, which could be due to the sensitivity limits of the Leica wide-field cryo-LM.

Although deconvolution of widefield cryo-LM has been successfully used for tissue imaging^36^, deconvolution has not been fully applied in cryo-LM of vitrified samples for cryo-CLEM applications. We show that the deconvolution of cryo-LM wide-field data is a necessary step for successful correlation and thus should be applied also to other cryo-CLEM strategies. Computationally performed tilt correction and slice extraction are critical to the workflow even for large organelles like LBs. When applied together, quantification showed that 76% of ABCA3-eGFP signal localized to membrane-bound structures and LB organelles. A residual 24% of signals could not be assigned to any MVB/LB-like structures, likely because of the limited axial resolution of widefield microscopy. Even though deconvolution can improve both lateral and axial resolution, it remains highly anisotropic and much larger than the 150-200 nm lamella thickness. The correlation precision of the described microscope setup is limited by the aberrations and the numerical aperture (NA) of 0.9 of the 50× objective used in the cryo-LM, which in particular affects resolution in the Z-direction. The development of lenses with improved aberration-correction, cryo-immersion lenses with a higher NA^37^ and the implementation of confocal or super-resolution cryo-LM can further improve the Z-resolution^38^ and thereby facilitate more accurate removal of out-of-lamella fluorescent signal in the post-correlation workflow. Overexpression of ABCA3-eGFP resulted in fewer LDs compared to naïve A549 cells (Supplementary Fig. 5). It is possible that the ABCA3 overexpression results in mobilization of neutral lipids in LDs for phospholipid synthesis. Thus, we were not able to use LDs as correlation markers and the final correlation was performed using the corners of the lamella which limited the X-Y correlation precision to 450 nm (Fig. 2k). In comparison, we were able to correlate fluorescently labelled LDs with a X-Y precision of 124 nm. Because of the characteristic spherical morphology of LDs and their increased electron density recognizable by cryo-EM, LDs can be used as fiducial markers to increase the correlation precision. Cryo-LM performed at low relative humidity (30 – 40%) improves both cryo-LM data and cryo-FIB milling. Therefore, integrating cryo-LM imaging into the cryo-FIB/SEM chambers, which allows obtaining fluorescence data before and after cryo-FIB milling^35^, is of great benefit to avoid sample transfers.

LB ultrastructure has long been subject to debate. Both parallel straight and concentric membrane sheets have been proposed as major structural components of LBs and both types were observed by CEMOVIS^18^. However, cutting artifacts and a lack of 3D data prevented a further understanding of the organelle architecture. Detailed analysis of membrane spacing revealed a repeat of 11 nm, compared to 7.3 nm found by CEMOVIS of rat lungs^18^. The difference in the membrane spacing could either be explained by the compression up to 50% that occurs during sectioning^19^, by maturation stage differences of the LBs, or by differences between human and rat LBs. The fact that primary human lung cells showed less compact packing of lamellar membranes than ABCA3 overexpressing A549 cells indicates that LBs membrane spacing is variable and the LB maturation might be dependent on ABCA3 levels and its regulation in a primary or immortalized cell line.

Post-correlation on-lamella cryo-CLEM combined with *in situ* cryo-ET of ABCA3-eGFP positive organelles allowed us to unravel LB architecture in 3D. The core exhibits multiple concentric membrane sheets with increasing curvature towards the center. However, none of the LBs observed were exactly centrosymmetric, which had previously been proposed based on polarized light microscopy^39^. “T”-junctions can be observed on the inner leaflet of the limiting membrane, where membranes are pushed into the lipidic core as parallel sheets consistent with a model proposed by Pérez-Gil^17^. In addition to parallel-curved membrane sheets, we occasionally observed crystalline lipid structures inside the LB core with a regular spacing of 3.4^-1^ nm as revealed by FFT. This spacing is in accordance with cholesterol ester crystals found in lipid droplets^40^.

Our finding that ABCA3-eGFP signal localizes not only to LBs but also to other membrane-bound organelles like MVBs, ER or vesicles is concurrent with previous data on trafficking and proteolytic processing of ABCA3 in LAMP3 positive vesicles^41^. However, ABCA3 distribution and its fraction on LB might be different in naive A549 cells and in primary lung cells. Since LBs found in ABCA3-eGFP expressing A549 cells do not only contain lamellated membranes but also crystalline lipidic structures and vesicles, these organelles rather represent ‘composite bodies’, a precursor form of LBs. LBs found in primary lung cells were morphologically similar to those induced by ABCA3-eGFP overexpression, indicating that neither the lamellar organization nor “T”-junctions found in the ABCA3-eGFP positive organelles are artifacts of overexpression.

Notably, we did not observe many vesicles inside LBs of HSAEpC cells, thus vesicles might not play a major role in LB biogenesis in primary lung cells. The A549 long-term culture showed an increased number of organelles compared to short-term non-transfected cells. Since no LB marker was used to localize LBs in A549 long-term culture, we performed cryo-ET on both cryo-FIB-milled cells and the periphery of whole cells. Interestingly, the majority of large LBs was found in the cell periphery of A549 long-term culture, where LB maturation and surfactant secretion occur. The post-correlation on-lamella cryo-CLEM workflow can be applied to study localization of other proteins involved in the maturation of LB such as Rab38, which plays an indispensable role in maintaining the morphology of LBs^42^.

We identified a dome-shaped protein complex on the LB outer membrane both in A549 cells and in primary lung cells that to our knowledge has not been reported yet. STA revealed that OMDPs exhibit a hollow cage-like structure, however, the limited number of the averaged OMDPs prevented obtaining molecular details of the complex. Nevertheless, based on the indicated C_8_ symmetry we speculate that the complex is built of multiple subunits composed of protein assemblies symmetrically organized around a single rotation axis. Notably, we frequently observed the structurally well-characterized 13 MDa major vault protein (MVP) in the cytosol of A549 cells, which forms large cage structures with 39-fold dihedral symmetry^43^. However, the dimensions of the terminal caps of the MVP are different from the OMDP, indicating that OMDP is not derived from MVP (Supplementary Fig. 9). Although OMDPs were rarely detected in ABCA3-eGFP expressing cells, they were frequently observed in A549 long-term culture. We believe that the low occurrence of OMDPs in ABAC3-eGFP overexpressing A549 cells can be explained by high ABCA3 occupancy on the limiting membrane, causing a displacement of other proteins. Since OMDPs have also been observed on MVBs, we assume that this complex plays a more general role in endo-lysosomal trafficking. The composition of the OMDP is beyond the scope of this study and warrants further investigation. Since mature LBs are exocytosed, we speculate that OMDPs could be involved in organelle tethering or in the regulation of fusion between the limiting membrane and the plasma membrane. An OMDP can be observed in a tomogram deposited by Ben Engel (EMDB 4604), indicating that the complex might be conserved across several domains of life and could be involved in membrane biology in general.

Based on structural analysis of ABCA3-positive organelles as well as LBs found in the A549 long-term culture, we propose a model for LB biogenesis (Fig. 6): ABCA3-mediated membrane lipid asymmetry causes the inner leaflet to push bilayers into the lumen forming “T”-junctions. It is conceivable that ABCA3 itself or other proteins are responsible for the concentration of phospholipids into membrane sheets. SP-B is a small protein that contains amphipathic helices^14^ and thus might be responsible for stabilizing the membrane sheets’ termini and preventing their fusion into vesicles. These sheets are detached from the LB limiting membrane and shaped according to the principal spherical curvature of the LB to maximize the lipid storage capacity per given volume. Finally, OMDP observed on the limiting membrane might regulate LB trafficking or be involved in exocytosis.

**Figure 6:**
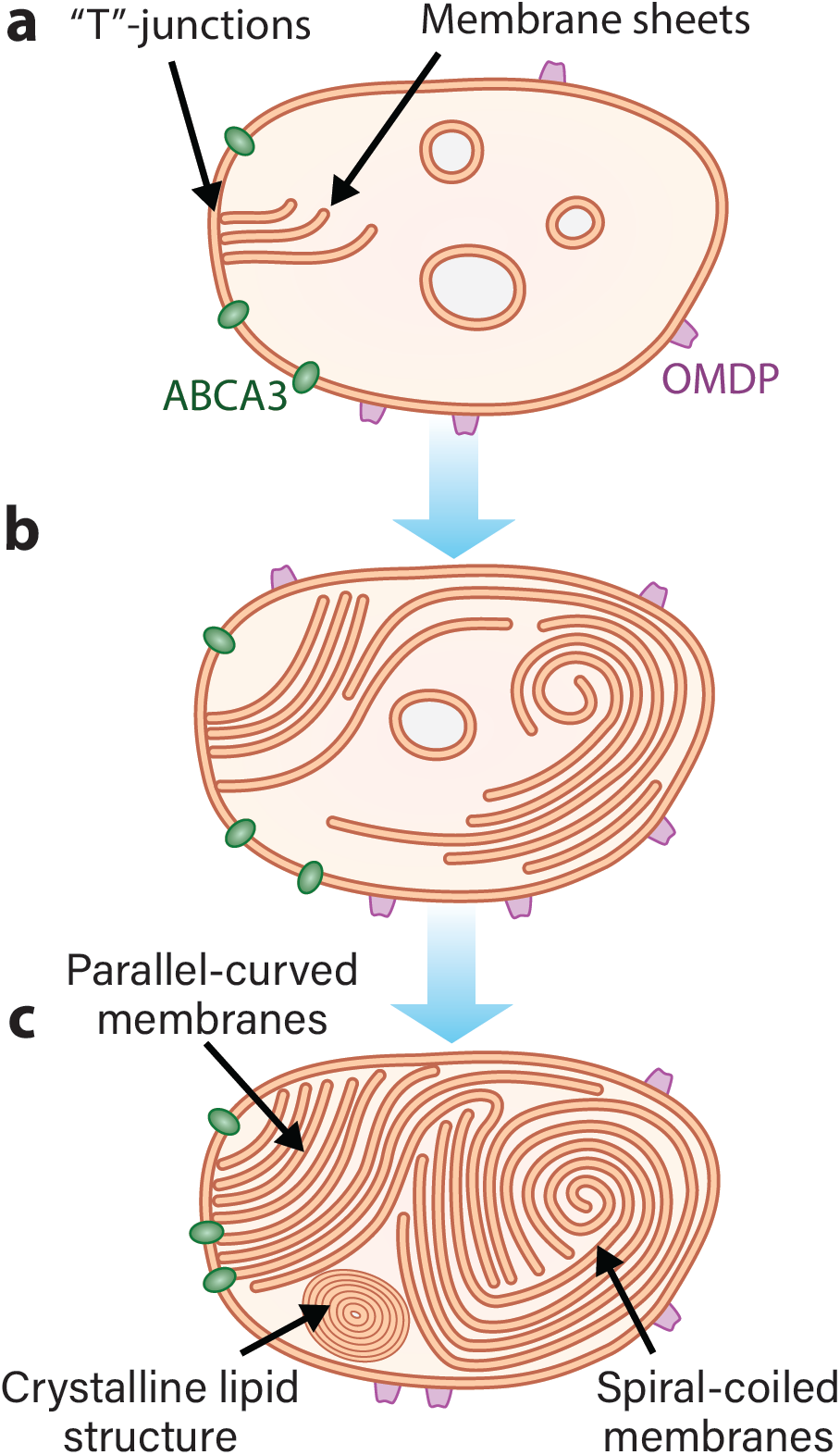
Proposed model of LB biogenesis. This model of LB biogenesis is based on observed ultrastructural features of LBs by cryo-TEM analysis. In all stages of LB biogenesis, OMDPs are present on the limiting LB membrane. **a**, Early stage: ABCA3 flips the phospholipids from the outer membrane leaflet to the inner one which results in a phospholipid asymmetry towards the inner leaflet. This leads to the formation of membrane sheets which are directly connected to the limiting membrane by “T”-junctions. **b**, Intermediate stage: membrane sheets grow further and are eventually detached from the limiting membrane. Individual membrane sheets start to organize in curved arrangements. **c**, Final stage: the lumen is packed with membrane sheets that form a combination of different structural arrangements including spiral-coiled, parallel straight and curved membranes as well as crystalline lipid structures.

In conclusion, we developed and implemented a robust post-correlation on-lamella cryo-CLEM workflow which can be applied to prove identity of cellular organelles using a well-defined fluorescent marker to reveal the structural details of organelles in the context of the native cellular environment.

## Methods

### Cell lines and cell culture

A549 cells^31^ were purchased from ECACC (Lot 16J012) and cultured in DMEM/F12 medium (ThermoFisher Scientific) supplemented with 10% (v/v) fetal bovine serum (FBS) (ThermoFisher Scientific) and 100 U/ml Penicillin-Streptomycin (ThermoFisher Scientific) at 37°C, 5% CO_2_. HSAEpC primary human lung cells were purchased from PromoCell (Lot 414Z012.2) and used at passage 3. The HSAEcP cells were maintained in Small Airway Epithelial Cell Basal Medium (PromoCell, C-21275) supplemented with Small Airway Epithelial Cell Growth Medium Supplement Mix (PromoCell, C-39175) and 100 U/ml Penicillin-Streptomycin (ThermoFisher Scientific) for 4 weeks (the medium was exchanged 3 times a week) and Detach Kit (PromoCell, C-41210) was used to detach the cells from the culture dish before seeding the cells on EM grids. A549 long-term culture was maintained without subculturing according to a protocol by Cooper *et al*.^*34*^ in Ham’s F12 (Sigma-Aldrich) supplemented with 10% FBS and 100 u/ml Penicillin-Streptomycin (ThermoFisher Scientific) for 28 days. The medium was exchanged twice a week. To prepare samples for plunge-freezing, 200 mesh Quantifoil™ Au R2/2 grids were plasma-cleaned in a Gatan Solarus 950 (Gatan) for 10 sec, sterilized by dipping into 70% Ethanol, placed in a 35 mm dish (ThermoFischer Scientific) coated with a thin layer of polydimethylsiloxane (PDMS, Dow Europe GmbH) and incubated in 1 ml of medium at 37°C for 30 min. Subsequently, the medium was removed and 1.2×10^5^ cells were seeded on grids placed in the PDMS-coated dish containing 2 ml of medium. The next day, cells were transiently transfected with 3 µg pE-hABCA3-eGFP plasmid (hABCA3-eGFP expression controlled by CMV promoter) in Trans-IT LT1 transfection reagent (Mirus). 48 h post-transfection the medium was removed, and the cells were incubated in fresh medium containing 1 µg/ml Hoechst 33342 (Sigma-Aldrich) and 150 nM LipiBlue (Dojindo Europe) to stain nucleus and lipid droplets, respectively. After 30 min incubation, cells were washed three times with medium.

### Vitrification

Cells were plunge-frozen into liquid ethane immediately after fiducial staining using a Leica EM GP2 automatic plunge-freezer. Cryogen temperature was set to -183°C and the chamber to 25°C and 95% humidity. Grids were blotted from the back with Whatman® Type 1 paper for 4 – 5 sec. 5 µl medium was added to the grid just before plunge-freezing. For samples that were used for whole-cell tomography, Protein-A Gold fiducials with a nominal diameter of 10 nm (Aurion) were added. Grids were inserted into AutoGrids™ (ThermoFisher Scientific) designed for FIB milling.

### Cryo-fluorescence widefield microscopy

Fluorescent maps of vitreous samples were acquired using a EM Cryo-CLEM widefield microscope (Leica) equipped with a 50× air objective (NA 0.9), Metal Halide light source (EL6000), an air-cooled detector (DFC9000GT) and a cryo-stage cooled to -190°C in the room with relative humidity ranging between 40 – 60%. A square of approximately 1.2 mm by 1.2 mm was acquired in the center of the grid using the LAS X Navigator. For each field of view, a symmetrical 30 µm Z-stack with a 300 nm spacing was acquired around the autofocus point. The following channels were used: TL-BF, DAPI (Ex: BP 350/50, Em: 460/50) and GFP (Ex: BP470/50, Em: 525/50). TIF files were exported from LAS X and a MIP map was stitched using the Grid/Collection stitching plugin^44^ in ImageJ/FIJI^45^.

### Deconvolution of cryo-fluorescence widefield microscopy data

To increase the clarity of the fluorescence signal, image stacks containing cells of interest were subjected to deconvolution in AutoQuant X3 (Media Cybernetics). A theoretical point-spread function (PSF) was utilized based on microscope and lens specifications (NA 0.9, 50× lens, refraction index 1) and refined over 100 iterations.

### Cryo-FIB-SEM milling

After cryo-fluorescence imaging, self-supporting lamellae were prepared using cryo-FIB milling on an Aquilos dual-beam FIB-SEM microscope in a room with controlled relative humidity between 30 – 40% (ThermoFisher Scientific) as first described by Rigort *et al*.^20^. Transfected cells were selected using correlation with a stitched MIP fluorescence map obtained in the previous step using the MAPS 3.3 software (ThermoFisher Scientific). Cells were coated with an organo-metallic platinum layer for 5 – 6 sec and gradually thinned in 4 steps at a stage angle of 15 – 18° using a Ga^+^ beam to yield lamellae with 150 – 200 nm thickness after the final milling step. If possible, micro-expansion joints were used as described by Wolff *et al*.^32^ to improve lamella stability. Progress was monitored by SEM imaging at 2.2 – 10 kV with ETD and T1 detectors.

### Tilt series acquisition and tomogram reconstruction

After lamella preparation, grids were transferred to a Titan Krios cryo-TEM operated at 300kV (ThermoFisher Scientific) equipped with a K3 direct electron detector and a Gatan imaging filter (Gatan). Image acquisition was controlled using SerialEM^46^. First, low-magnification maps were acquired at 135× magnification to find the lamellae, then medium-magnification maps (MMM) were acquired at 8,700× for correlation and the identification of sites for tilt series (TS) acquisition at 26,000× or 33,000× (corresponding pixel sizes at the specimen level: 3.356 and 2.671 Å, respectively). TS were acquired using a dose-symmetric scheme^47^ with a constant electron dose of approx. 3 e^-^/Å^2^ per projection, target defocus -5 μm, energy filter slit at 20 eV, covering the range from 60° to -60° in 3° increments. On-lamella tomography was done with a stage tilted to 6° to compensate for the pre-tilt of the lamella with respect to the grid. Each projection was acquired in counting mode using dose fractionation and 20 – 40 individual frames, which were aligned and summed using the SEMCCD plugin in SerialEM. Whole-cell tomography was performed as stated above with the exception that the TS were acquired at magnification 33,000× (pixel size at the specimen level: 4.302 Å) using a K2 direct electron detector instead of K3 direct electron detector.

TS were processed using the IMOD package^47^. TS were aligned using patch-tracking. Before reconstruction, the contrast transfer function (CTF) was corrected by phase-flipping and the micrographs were dose-filtered. Tomograms were reconstructed using the weighted back-projection algorithm with a SIRT-like filter equivalent to 10 iterations. For power spectral analysis in Fig. 4, tomograms were reconstructed using the SIRT algorithm implemented in 3dmod.

### 3D rendering of reconstructed tomograms

Renderings were created in Amira 2019.3 (ThermoFisher Scientific). First, a Membrane Enhancement Filter was applied. Using the Top Hat segmentation tool, a first 3D rendering was created. Based on this initial model, all membranes were manually segmented.

### Subtomogram averaging

STA was performed in Dynamo^33^. Particles (n = 105) were picked manually in 2 tomograms and 100 voxel subtomograms were extracted using a crop-on-wall function with the initial particle orientation assigned normal to the segmented limiting membrane of the LB. An initial reference was generated by averaging all 105 particles and the average was iteratively refined using a half-dome shaped mask. A symmetry scan was performed using a focused mask on the top of the cage assembly. Subsequently, the average was further refined using the C_8_ symmetry.

### Post-correlation on-lamella correlation workflow

After vitrification of fluorescently labeled cells by plunge freezing, grids were mapped on the Leica cryo-LM. The grid was transferred to a dual-beam cryo-FIB/SEM. By low-precision correlation of the MIP fluorescent map with the SEM map of the grid, cells of interest were selected and FIB-milled. After a second transfer of the sample to the 300 kV cryo-TEM, the lamellae were mapped, areas of interest were selected, and TS were acquired. To facilitate high precision post-correlation of the lamellae, the grids were recovered after the cryo-ET acquisition and the lamellae were mapped a second time in the cryo-LM to reveal the exact lamella positions on the grid. To merge the fluorescent signal from the first cryo-LM map with the information of the lamella position on the second cryo-LM map, the deconvolved stacks were registered in MATLAB 2018b (MathWorks) using the *imregtform* function with a custom script (see code availability). A rigid transformation matrix including translation and rotation was calculated based on the same fluorescent channel of both cryo-LM maps respectively. The remaining channels including TL-BF from the second cryo-LM map were transformed by applying the calculated transformation matrix. The fluorescent channels of the first cryo-LM map were merged with the transformed TL-BF channel of the second cryo-LM in FIJI/ImageJ and saved as a composite TIF. To compensate for the FIB-milling angle, the composite stack was resliced using FIJI/ImageJ. Using the TL-BF channel, the actual lamella tilt in the stack was measured, and the composite stack was rotated accordingly to compensate for the tilt. The Z-slice of the lamella was identified and extracted after a second reslice in FIJI/ImageJ. This single tilt-corrected slice was correlated with the stitched TEM-map using a non-rigid 2D transformation in ec-CLEM^48^ utilizing the four corners of the lamella as well as clearly identifiable fluorescently labeled organelles like lipid droplets as landmarks.

### Correlation precision analysis

To evaluate the correlation and the X-Y precision of the post-correlation on-lamella cryo-CLEM workflow, the final correlation of the MMM and cryo-LM map was analyzed. Local maxima of the cryo-LM map were calculated using FIJI/ImageJ using a tolerance threshold of 10. For each local maximum, ultrastructures of the correlated MMM in a spherical area (radius = 1 µm) were classified either as LD, LB or membrane-bound organelle (MVBs, ER, vesicle). Mitochondrial or nuclear membranes as well as areas without any membrane-bound structures were classified as unspecific. For each assigned ultrastructure, the distance and angle to the center of the local maximum was measured and plotted as a radar plot to estimate the correlation precision.

### Confocal fluorescence microscopy

Confocal microscopy of A549 cells transfected with ABCA3-eGFP was done using a SP8 TCS laser scanning confocal microscope (Leica) equipped with a 63× 1.4 NA oil immersion objective and an environmental control chamber heated to 37°C. Image analysis was performed in ImageJ/FIJI^45^. Automated 3D segmentation of ABCA3-eGFP structures was done in Imaris 9.5.1 (Bitplane) after deconvolution in AutoQuant X3.1 (Media Cybernetics).

### Code availability

The FIJI/ImageJ macro for fluorescent map assembly and the MATLAB code used for 3D stack registration are. available on github under https://github.com/chlanda-lab.

## Acknowledgments

We would like to thank Dr. Surafel Mulugeta for kindly providing the plasmid hABCA3-eGFP and to Prof. Dr. Hans-Georg Kräusslich for providing the HSAEpC cells. We thank Dr. Stefan Pfeffer for critical reading of the manuscript and to Dr. Ben Engel and Dr. Lars Anders Carlson for a fruitful discussion about OMDP. We would like to acknowledge microscopy support from the Infectious Diseases Imaging Platform (IDIP) at the Center for Integrative Infectious Disease Research Heidelberg. We would like to acknowledge access to the infrastructure and support provided by the Cryo-EM Network at the Heidelberg University (HD-cryoNet). Funded by the Deutsche Forschungsgemeinschaft (DFG, German Research Foundation) Projektnummer 240245660 SFB 1129.

