## Supplementary data for "Post-correlation on-lamella cryo-CLEM reveals the membrane architecture of lamellar bodies"

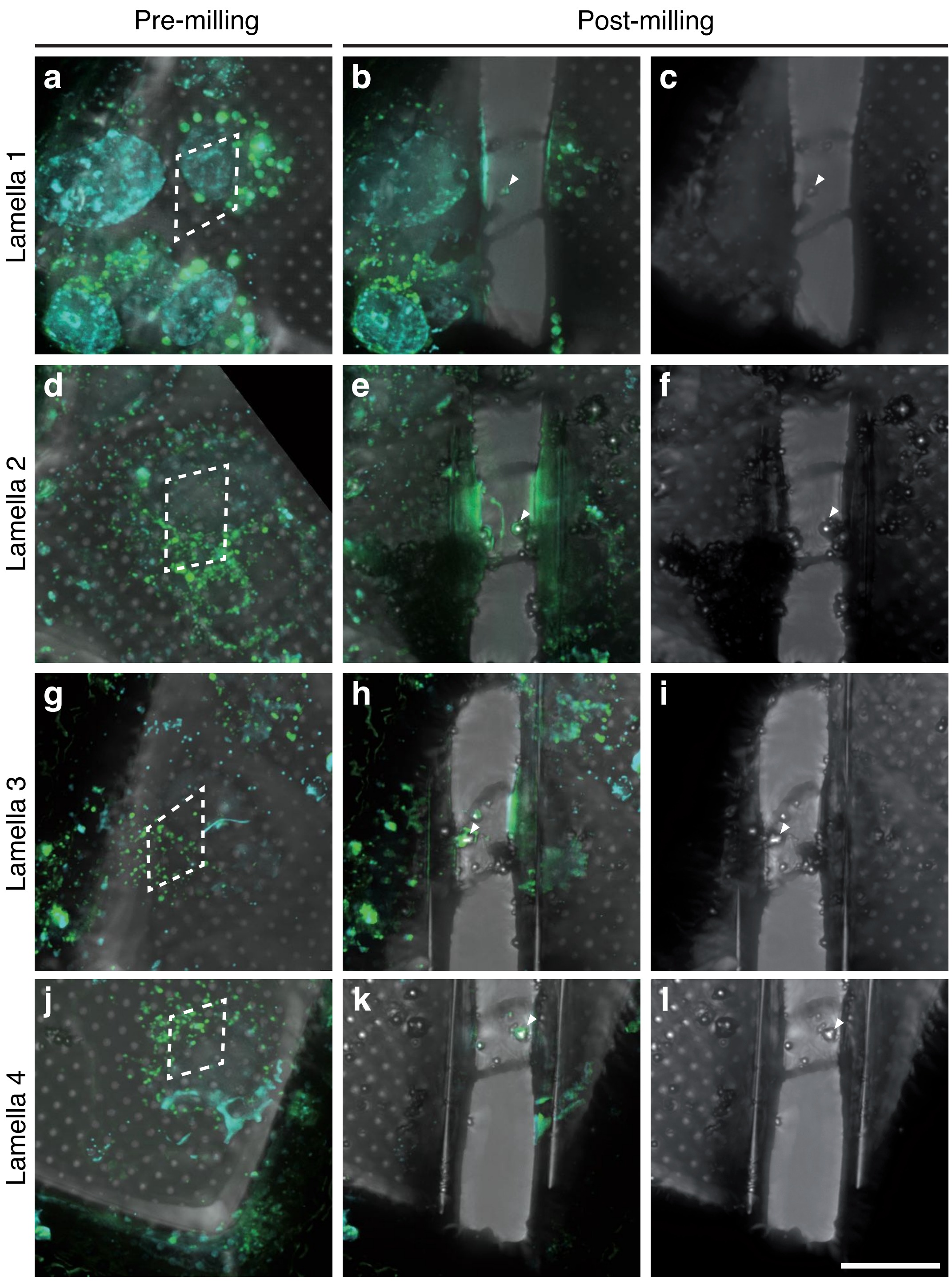


**Supplementary Fig. 1: Fluorescence signal is lost on lamella sites after cryo-FIB milling and cryo-ET acquisition.** All panels show maximum intensity projections of a cryo-LM stack with a 300 nm Z-spacing. **a, d, g, j,** Cryo-LM image acquisition of pre-milled grids show a preserved fluorescence signal of ABCA3-eGFP (green), nucleus and LD staining (cyan) for 4 different lamellae. **b, e, h, k,** After cryo-FIB milling and cryo-ET image acquisition of the lamellae, the fluorescence signal of ABCA3-eGFP, nucleus and LD staining are completely lost on all lamellae, although fluorescence signal of the surrounding cell body is preserved. The lamella sites are indicated as dotted outlines in the pre-milled state (a, d, g, j). **c, f, i, l,** TL-BF channel of the lamella sites show deposition of ice contamination on the lamellae (arrowhead). These contaminations lead to an unspecific autofluorescence signal present in all fluorescent channels (arrowheads in b, e, h, k). Scale bar: 20 µm.

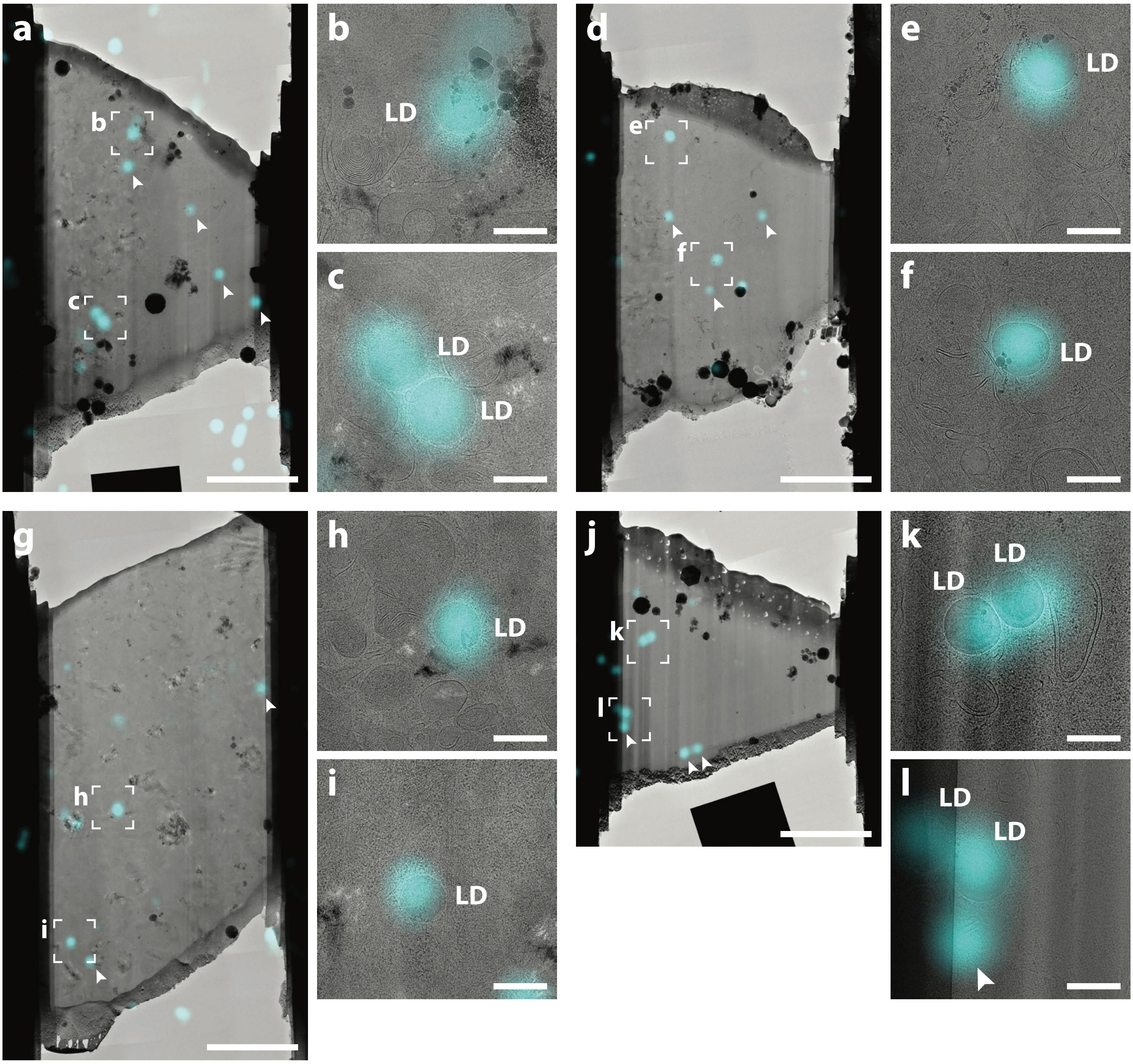


**Supplementary Fig. 2: Benchmark of post correlation on-lamella cryo-CLEM using fluorescently labelled lipid droplets.** A549 cells were fluorescently labelled with a neutral lipid dye (LipiBlue), shown in cyan, to stain lipid droplets (LD). After plunge freezing, the correlation workflow was applied on 4 lamellae (a, d, g, j). All LDs (n = 11) which were present on the 4 lamellae colocalize with the fluorescent signal. Areas of correlated LDs are indicated as squares on the overview TEM maps of the lamellae and are shown with a higher magnification in (b, c, e, f, h, i, k, l). Out-of-lamella fluorescent signals (n = 12), which do not correlate with LDs in the TEM image are marked with arrowheads. Scale bars: (a, d, g, j) 5 µm, (d, c, e, f, h, i, k, l) 500 nm.

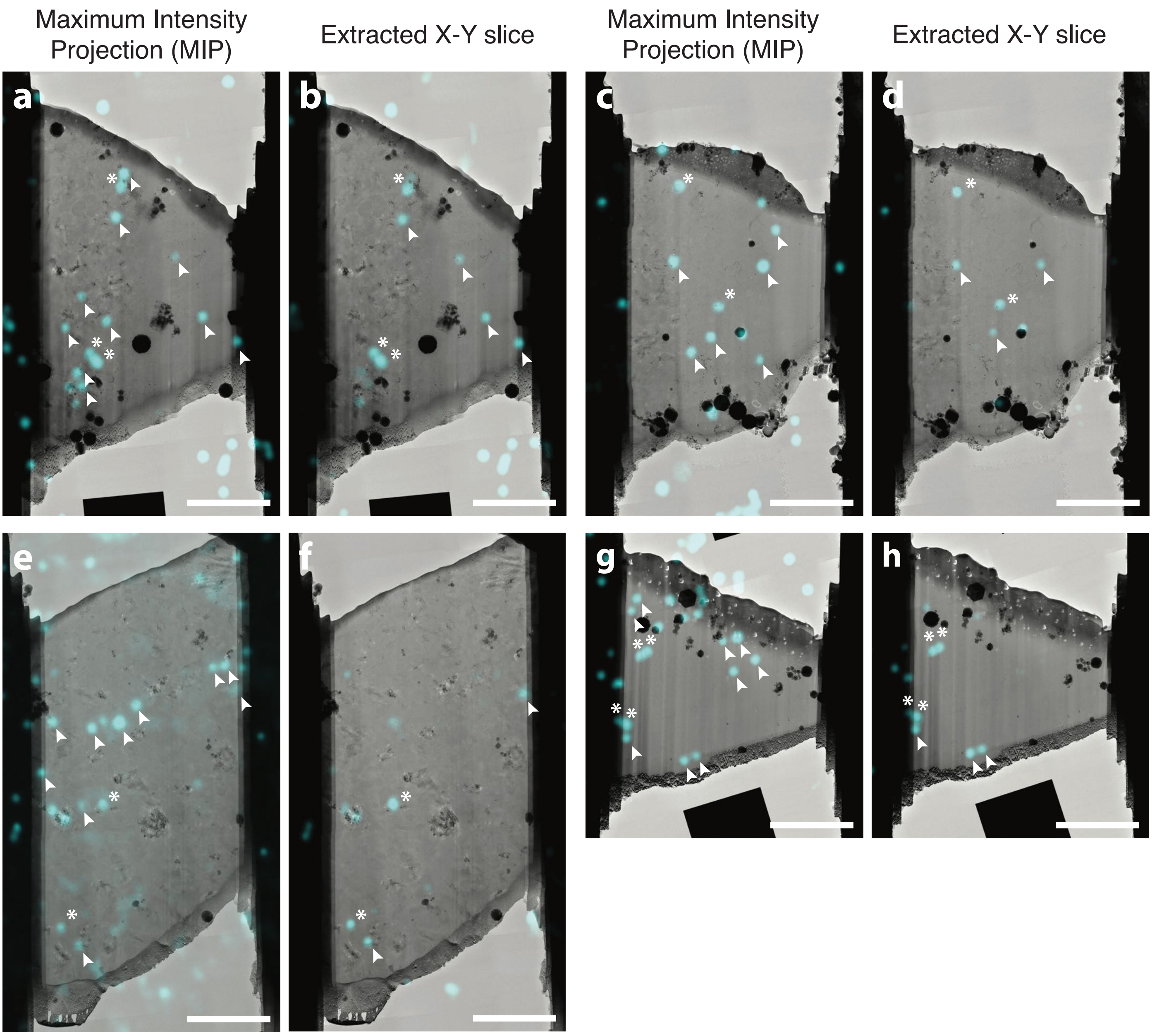


**Supplementary Fig. 3: Comparison of correlation results using either a MIP or an extracted X-Y slice of the fluorescent stack.** LDs of A549 cells were fluorescently labelled (LipiBlue) and the post-correlation on-lamella cryoCLEM workflow was conducted. In order to analyze the impact of the tilt correction and the extraction of a single X-Y slice, the correlation was performed using a MIP projection of the fluorescent stack (a, c, e, g) or a single extracted slice (b, d, e, g) on the 4 lamellae shown in Supplementary Fig. 2. All LDs (n = 11) present in the 4 lamellae (asterisks) correlated to the fluorescent signal. Out-of-lamella signals (arrowheads) which correspond to LDs removed during cryo-FIB milling. The number of out-of-lamella signals is reduced by 66% (from 35 to 12) when using a single extracted slice in comparison to the correlation using a MIP. Scale bars: 5 µm.

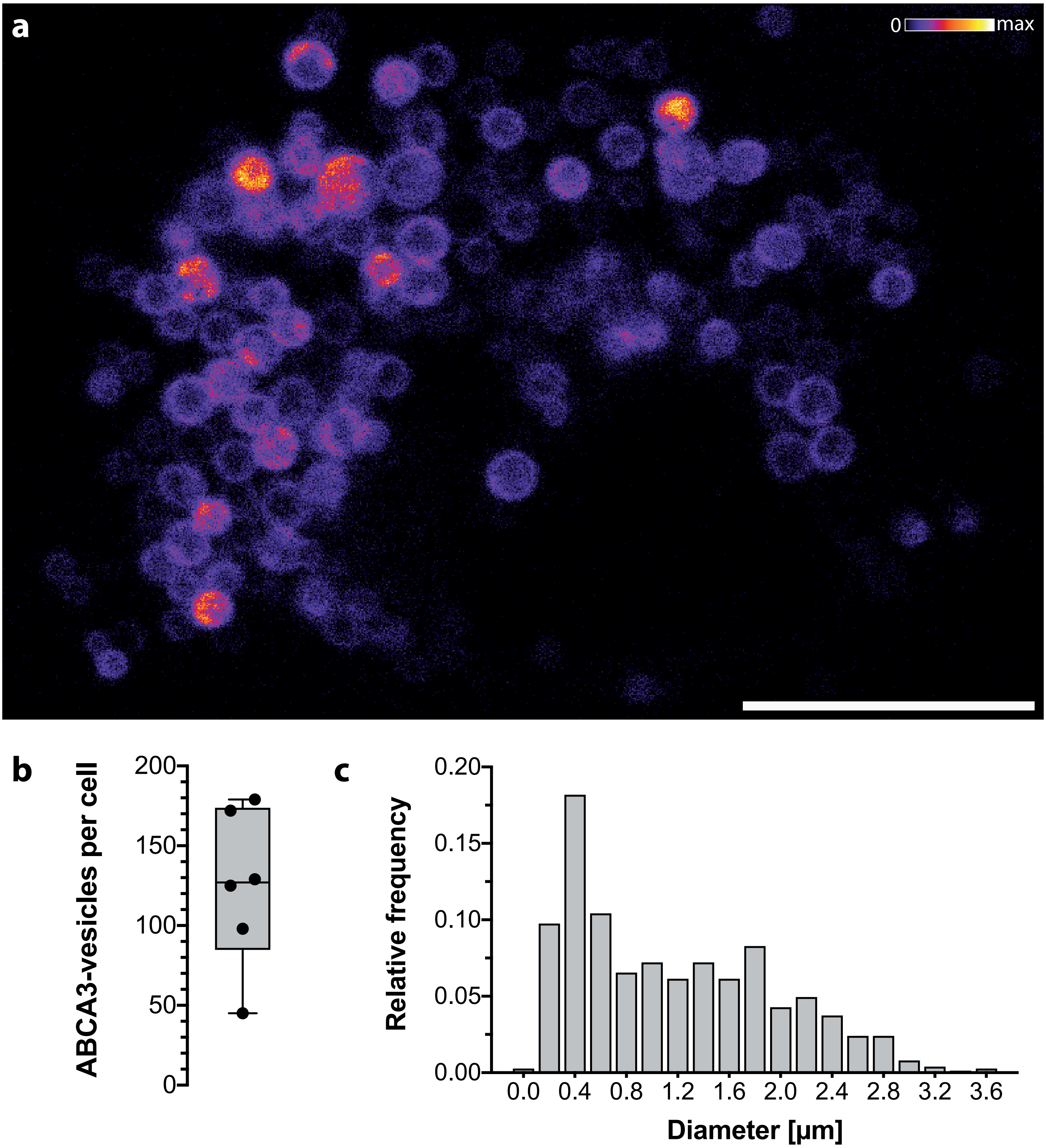


**Supplementary Fig. 4: Analysis of ABCA3-eGFP positive vesicles in transiently transfected A549 cells. a**, Confocal image slice of a live A549 cell expressing ABCA3-eGFP for 48h. **b**, Confocal volume data of 6 individual cells were semi-automatically 3D segmented using Imaris (Bitplane). The number of ABCA3-eGFP positive vesicles per cell varies considerably between cells. Box and whiskers blot indicating all data points, median (127), minimum (45), maximum (179) as well as the 25% quartile (85) and the 75% quartile (174). **c**, The outer diameter of all segmented ABCA3-eGFP positive vesicles of the 6 cells were analyzed as a histogram with bin size of 0.2 µm, which reveals a mean diameter of 1.2 µm (SD 0.8 µm) with a range between 0.1 - 4.5 µm. Scale bar: 10 µm.

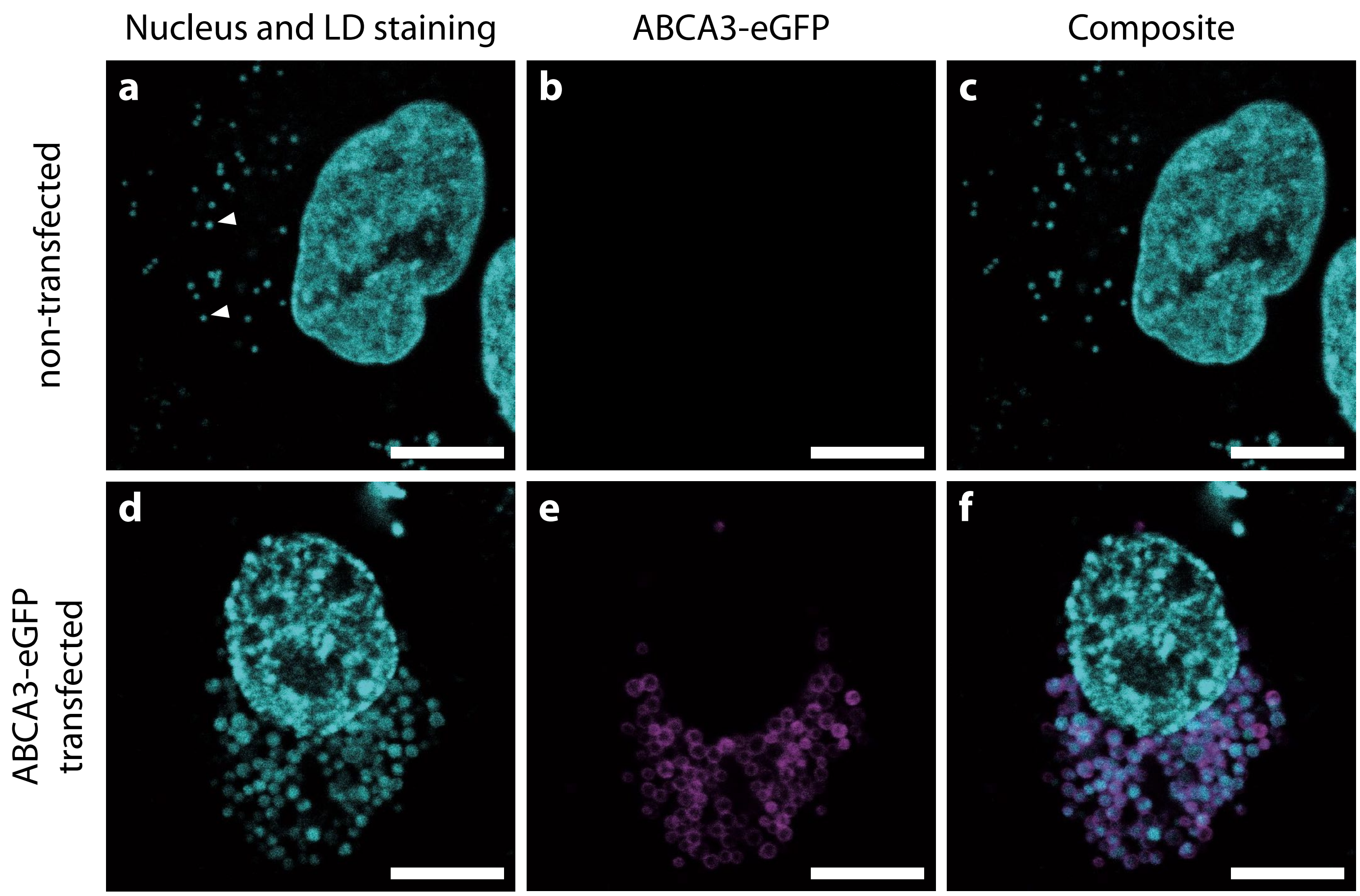


**Supplementary Fig. 5: ABCA3-eGFP positive organelles colocalize with neutral lipid dye.** A549 cells were transfected with ABCA3-eGFP. Nuclei (Hoechst 33342) and neutral lipids (LipiBlue) were fluorescently labeled and live-cell confocal image stacks were acquired. Each image represents one extracted confocal image slice with nucleus and neutral lipids in cyan and ABCA3-eGFP in magenta. **a–c**, The neutral lipid signal in non-transfected A549 cells reveals a lipid droplet morphology. Two exemplary lipid droplets are marked with an arrowhead. **d–f**, In contrast to non-transfected cells, A549 cells transfected with ABCA3-eGFP feature a different morphology: organelles labelled with neutral lipid dye show a larger diameter and a weaker signal intensity. Furthermore, the neutral lipid signal colocalizes with the ABCA3-eGFP signal. Scale bars: 10 µm.

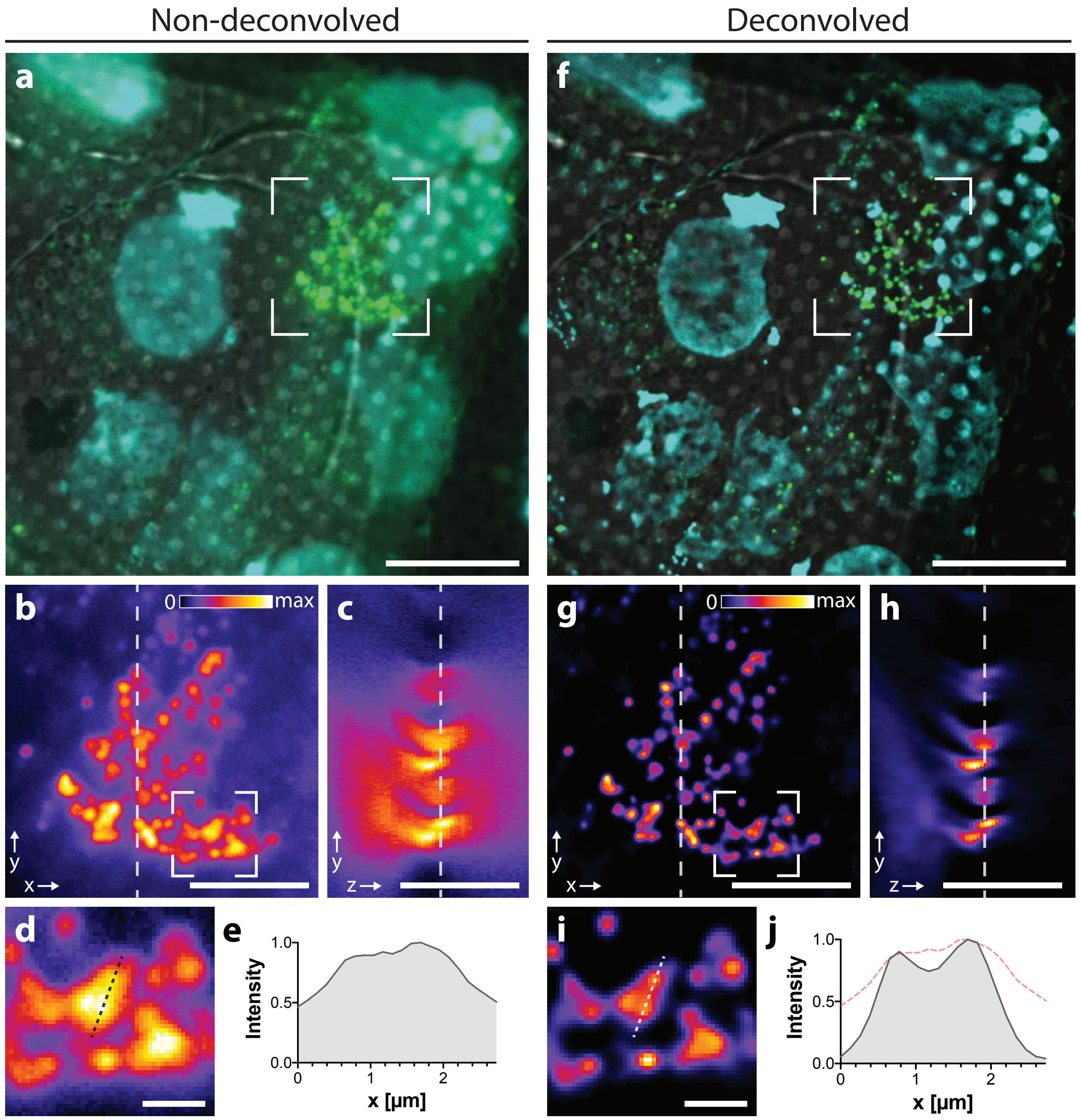


**Supplementary Fig. 6: Deconvolution improves resolution of cryo-LM wide-field fluorescent images.** A549 cells were grown on EM-grids and transfected with ABCA3-eGFP. Nuclei and lipid droplets were fluorescently labeled. After vitrification, the EM-grid was mapped by cryo-LM, acquiring 30 µm thick Z-stacks with a spacing of 300 nm. (a–e) show the data before deconvolution, and (f–j) after deconvolution. **a**, **f,** Single XY-slice of the acquired Z-stack with the TL-BF channel in grayscale, ABCA3-eGFP in green and the combined signal of the lipid droplet and nucleus stain in cyan. The images show the full dynamic range of the data. **b** and **g**, Details of the same area showing only the ABCA3-eGFP signal, while **c** and **h** show a single ZY plane. Respective plane positions are indicated by dashed lines. **d** and **i**, For further analysis, areas indicated in (b) and (g), are enlarged and intensity was measured along the indicated lines. **e**, Line profile of adjacent LBs before deconvolution. **j**, Line profile showing separation of signals after deconvolution (gray). Profile before deconvolution is given as comparison (dashed red line). Scale bars: (a and f) 20 µm, (b, c, g, h) 10 µm, (d & i) 2 µm.

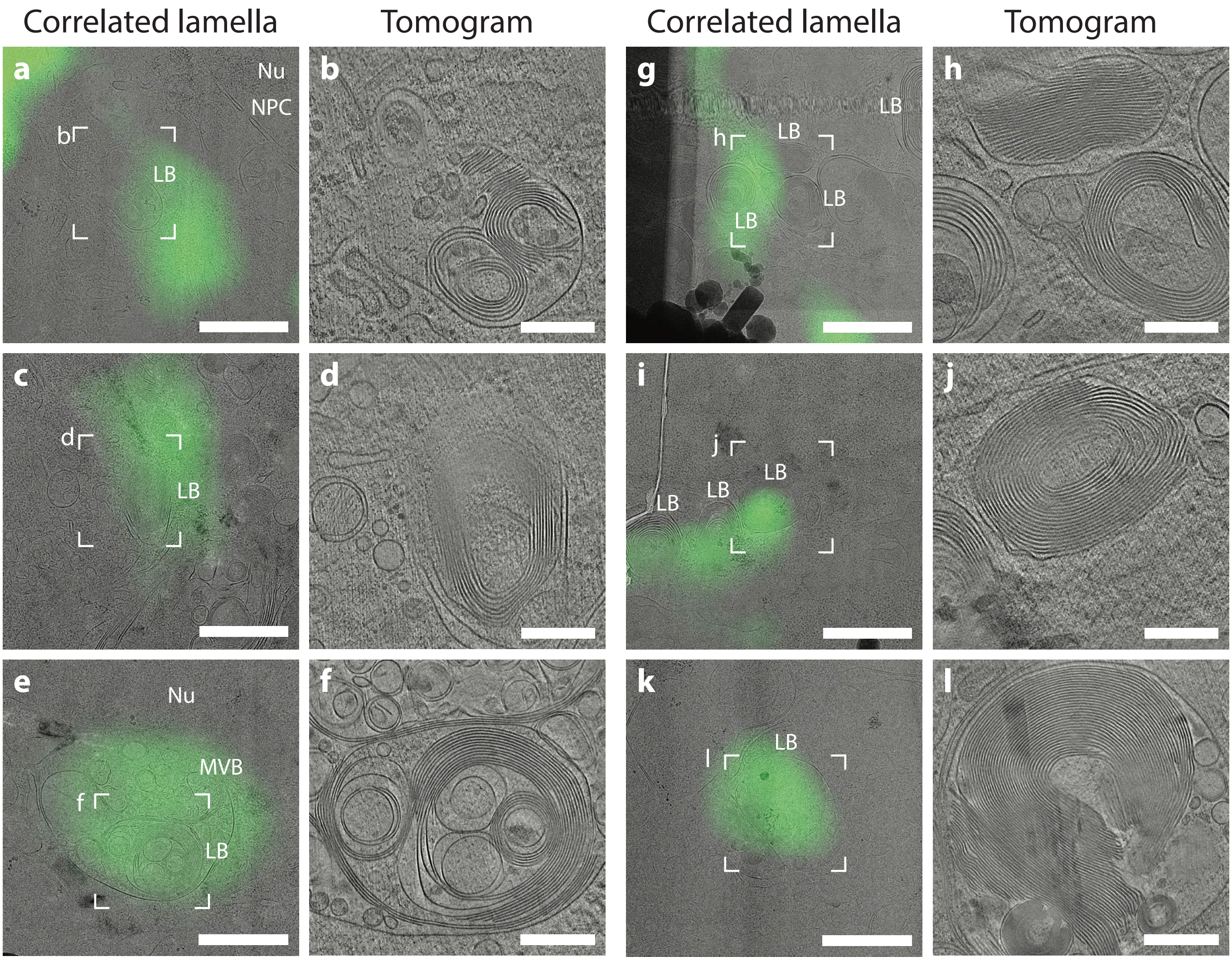


**Supplementary Fig. 7: Overview of post-correlation on-lamella cryo-CLEM data of A549 cells overexpressing ABCA3-eGFP.** A549 cells were transfected with ABCA3-eGFP and processed by the post-correlation on-lamella cryo-CLEM workflow. Correlated lamellae used in this study are shown as correlated overviews of the stitched MMM with the cryo-LM map. Tomogram acquisition sites are marked as white squared and a central slice of the reconstructed tomograms are shown. Scale bars: (a, c, e, g, i, k) 1 µm, (b, d, f, h, j, l) 300 nm.

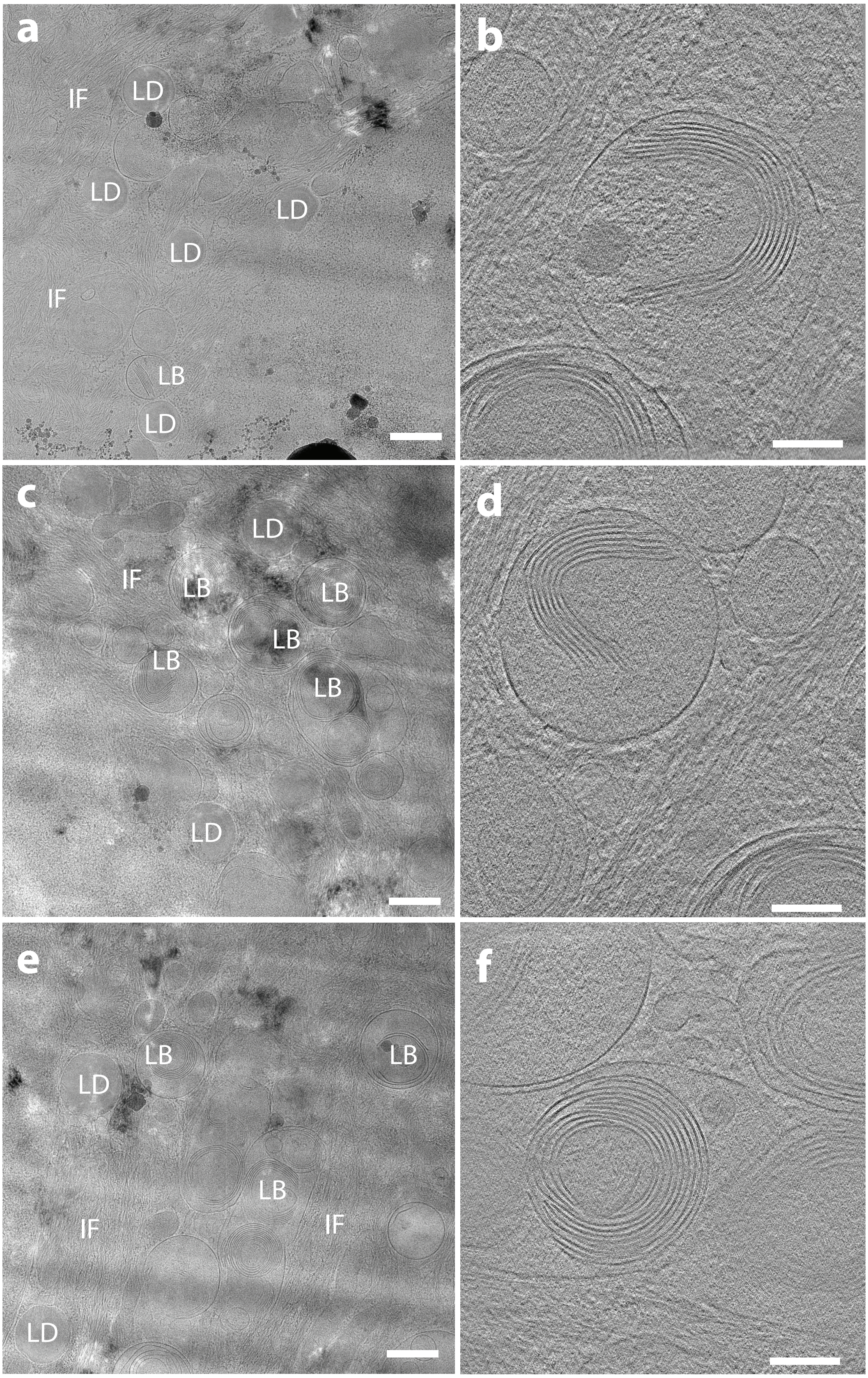


**Supplementary Fig. 8: *In situ* cryo-ET of cryo-FIB milled HSAEpC cells.** HSAEpC cells were seeded on EM grids after 4 weeks culturing without splitting. **a, c, e** Cryo-EM images of cryo-lamellae showing areas with multiple lamellar bodies (LB), intermediate filaments (IF) and lipid droplets (LD). **b, d, f,** Slices of tomograms capturing LBs. Scale bars: (a, c, e) 500 nm, (b, d, f) 200 nm.

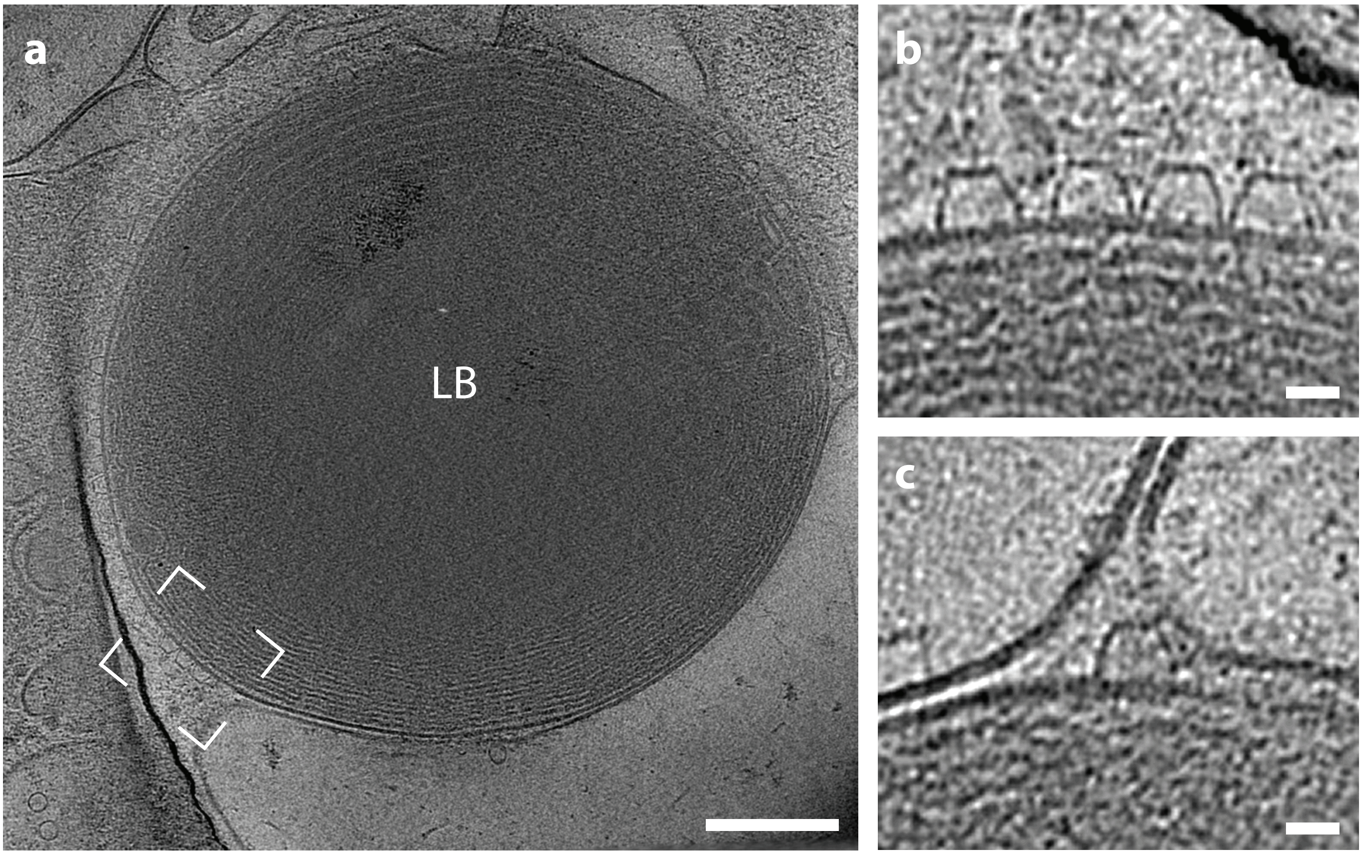


**Supplementary Fig. 9: Slice of a tomogram showing a large LB at the cell periphery of long-term cultured A549 cells. a,** Cells cultured on EM grids were imaged whole-cell cryo-ET. **b,** Detail of a tomogram marked in (a) showing OMDPs. **c,** Detail of a tomogram corresponding to a marked region in (a) showing OMDP in proximity to a membrane of a neighboring organelle. Scale bars (a) 200 nm, (b, c) 20 nm.

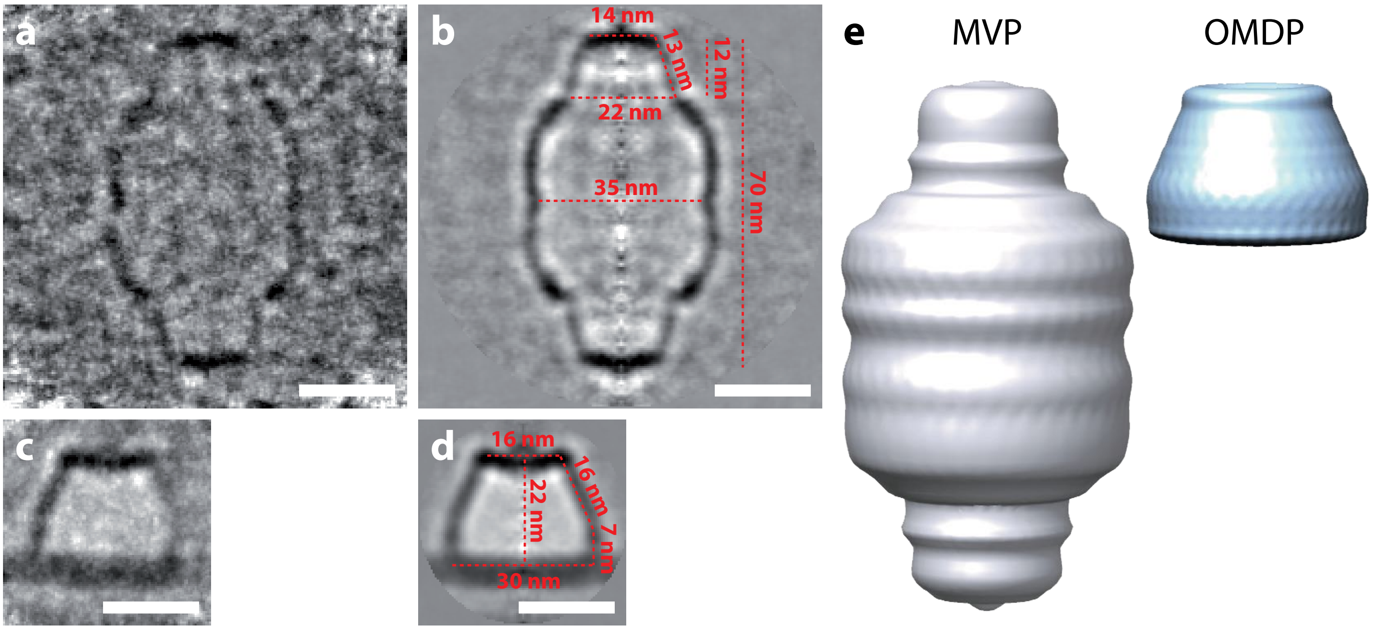


**Supplementary Fig. 10: Side-by-side comparison of MVP and OMDP complexes.** Subtomograms containing MVP (n=25) and OMDP (n=105) were manually extracted, aligned and averaged using Dynamo. Symmetry C39 was applied to both MVP and OMDP during the alignment. **a, b**, Central slices of the MVP non-symmetrized and symmetrized average, respectively. **c, d**, Central slices of the OMDP non-symmetrized and symmetrized average, respectively. Dimensions of the complexes are marked by a red dashed line. **e**, Isosurfaces of symmetrized averages of the MVP and OMDP complexes at the same scale. Scale bars: 20 nm.
